# N-Terminomic Changes of Neurons During Excitotoxicity Reveal Proteolytic Events Associated with Synaptic Dysfunctions and Inform Potential Targets for Neuroprotection

**DOI:** 10.1101/2022.03.13.484119

**Authors:** S. Sadia Ameen, Nane Griem-Krey, Antoine Dufour, M. Iqbal Hossain, Ashfaqul Hoque, Sharelle Sturgeon, Harshal Nandurkar, Dominik F. Draxler, Robert L. Medcalf, Mohd Aizuddin Kamaruddin, Isabelle S. Lucet, Michael G. Leeming, Dazhi Liu, Amardeep Dhillon, Jet Phey Lim, Faiza Basheer, Hong-Jian Zhu, Laita Bokhari, Carli Roulston, Prasad N. Paradkar, Oded Kleifeld, Andrew N. Clarkson, Petrine Wellendorph, Ciccotosto D. Giuseppe, Nicholas A. Williamson, Ching-Seng Ang, Heung-Chin Cheng

## Abstract

Excitotoxicity is a neuronal death process initiated by over-stimulation of ionotropic glutamate receptors. Although dysregulation of proteolytic signaling networks is critical for excitotoxicity, the identity of affected proteins and mechanisms by which they induce neuronal cell death remain unclear. To address this, we used quantitative N-terminomics to identify proteins modified by proteolysis in neurons undergoing excitotoxic cell death. We found that most proteolytically processed proteins in excitotoxic neurons are likely substrates of calpains, including key synaptic regulatory proteins such as CRMP2, doublecortin-like kinase I, Src tyrosine kinase and calmodulin-dependent protein kinase IIβ (CaMKIIβ). Critically, calpain-catalyzed proteolytic processing of these proteins generates stable truncated fragments with altered activities that potentially contribute to neuronal death by perturbation of synaptic organization and function. Blocking calpain-mediated proteolysis of one of these proteins, Src protected against neuronal loss in a rat model of neurotoxicity. Extrapolation of our N-terminomic results led to the discovery that CaMKIIα, an isoform of CaMKIIβ undergoes differential processing in mouse brains under physiological conditions and during ischemic stroke. In summary, our findings inform excitotoxic neuronal death mechanism and suggest potential therapeutic strategies for neuroprotection.

**In Brief:** Ameen, et al. used a proteomic method called N-terminomics to identify proteolytic events occurring in neurons during excitotoxicity. They found that most proteolytic processing is mediated by calpains, resulting in the generation of stable truncated fragments with the potential to induce synaptic dysfunction and loss, eventually leading to neuronal death. They further showed that some of these proteolytic processed proteins, such as the protein kinases Src and CaMKII, are potential targets for neuroprotection.

**Highlights:**

- Identification of over 300 neuronal proteins cleaved by calpains to form stable truncated fragments during excitotoxicity.
- The calpain cleavage sites of these proteins unveil for the first time the preferred cleavage sequences of calpains in neurons.
- These pathological proteolytic events potentially induce synaptic dysfunction and loss, which likely contribute to excitotoxic neuronal death.
- Some of the neuronal proteins proteolyzed by calpains are potential targets of neuroprotection.

**Graphical abstract: Pathological proteolytic events in neurons during excitotoxicity unveiled by N-terminomic analyses:** **(A)** N-terminomic and global proteomic analyses identified neo-N-terminal sites and neuronal proteins undergoing significant abundance changes during excitotoxicity. **(B)** Informatic analysis of the proteomic results predicted (i) the preferred sequences of proteolytic processing of neuronal proteins catalyzed by calpains during excitotoxicity and (ii) perturbation of synaptic organization and functions as the major consequence of calpain-mediated proteolytic events. **(C)** Validation of these predictions and further experimentations unveiled: (i) calpain-mediated cleavage of proteins associated with synaptic damage in excitotoxic neurons, (ii) a new mechanism of dysregulation of CaMKIIα and CaMKIIβ, which are key protein kinases governing synaptic dysfunctions and excitotoxic neuronal death and (iii) potential therapeutic targets such as the protein kinases Src and CaMKII for neuroprotection

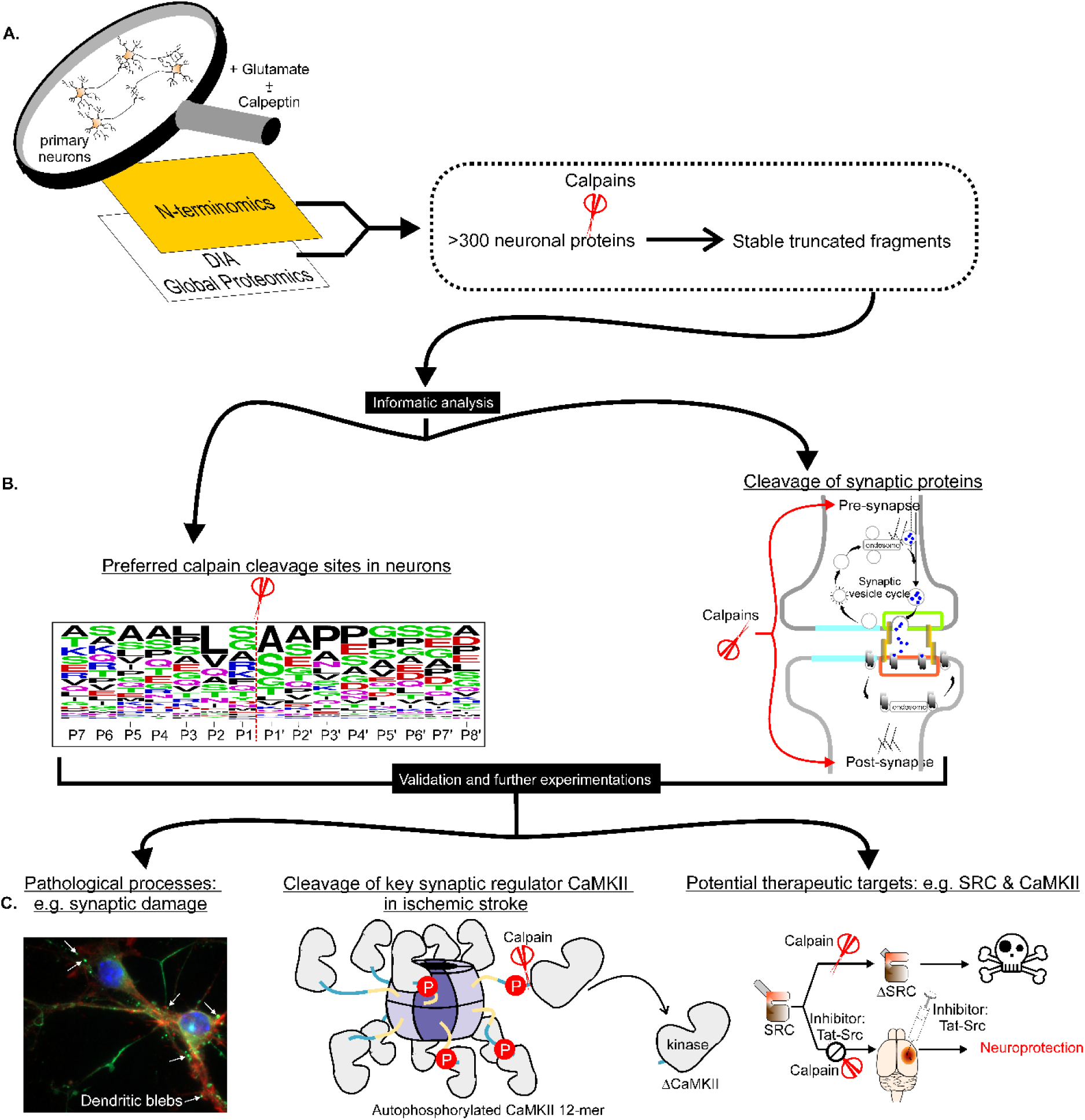

**One Sentence Summary:** Proteolytic events in neurons during excitotoxicity inform neuronal death mechanism and potential therapeutic strategies for neuroprotection.

## INTRODUCTION

Excitotoxicity is a pathological cell death process that underpins neuronal cell loss in multiple acute and chronic neurological disorders such as ischemic stroke, Parkinson’s and Alzheimer’s diseases (reviewed in (Fricker, Tolkovsky et al., 2018)). There are no FDA-approved pharmacological agents to protect against excitotoxic neuronal loss in neurological disorders even after decades of intensive studies of the molecular mechanisms of neuronal cell death (Chamorro, Lo et al., 2021, Savitz & Fisher, 2007). However, recent findings in a phase III stroke trial have highlighted putative neuroprotective compounds targeting pathologically activated signaling events directing excitotoxic neuronal death as a feasible therapeutic option (Hill, Goyal et al., 2020, Lipton, 2007). As such, further investigations to define the mechanisms of excitotoxic neuronal death with the aim to uncover the pathologically activated signaling events as therapeutic targets are critical to development of these neuroprotective compounds (reviewed in (Ballarin & Tymianski, 2018, Fisher & Savitz, 2022)).

Excitotoxicity is initiated by over-stimulation of ionotropic glutamate receptors (iGluRs), especially N-methyl-D-aspartate (NMDA) receptors (Choi, 1988, Olney, 1969, Simon, Swan et al., 1984), which permit excessive influx of extracellular calcium (Ca^2+^) into the cytosol to hyperactivate proteases (Ginet, Spiehlmann et al., 2014, Lankiewicz, Marc Luetjens et al., 2000, Wang, Nath et al., 1996, Yamashima, Kohda et al., 1998), neuronal nitric oxide synthase (nNOS) (Sattler, Xiong et al., 1999) and NADPH oxidase 2 (NOX2) (Brennan, Suh et al., 2009). The excitotoxicity-activated proteases cleave specific neuronal proteins to dysregulate their activities, biological functions and stability (Tominaga, Nakanishi et al., 1998, Wang et al., 1996), thereby contributing to neuronal cell death. Amongst the neuronal proteins dysregulated by these proteases are protein kinases and phosphatases, whose proteolytic processing can contribute to neuronal cell death by altering the phosphorylation states of specific neuronal proteins (Hossain, Roulston et al., 2013, Meyer, Torres-Altoro et al., 2014, Shioda, Moriguchi et al., 2006, Wang, Liu et al., 2003, Wu, Tomizawa et al., 2004). Identification of the proteases, protein kinases and phosphatases activated during excitotoxicity and their substrates in neurons are thus critical for charting the signaling pathways and pathologically-induced cellular events governing excitotoxic neuronal death, and for discovering novel neuroprotective therapeutic approaches (Lipton, 2007).

We previously used mouse cortical neurons, a well-defined *in vitro* model of excitotoxicity to profile excitotoxicity-related changes in phosphorylation of neuronal proteins (Hoque, Williamson et al., 2019). To gain further insight into post-translational modification events involved in excitotoxic neuronal death, the present study sought to identify the substrates of the excitotoxicity-activated proteases and understand how proteolysis alters their biological functions. To do this, we used a quantitative N-terminomics procedure called Terminal Amine Isotopic Labelling of Substrates (TAILS) (Kleifeld, Doucet et al., 2010) to identify and quantify the N-terminal peptides derived from N-termini of cellular proteins in control neurons and neurons undergoing excitotoxic cell death. Results of our TAILS analysis document changes in stability and the N-termini of neuronal proteins proteolyzed by activated proteases during excitotoxicity induced by glutamate over-stimulation. Herein we describe results of our analysis and illustrate how the results unveil new mechanisms by which the excitotoxicity activated proteases dysregulate key proteins controlling synaptic organization and functions. Some of these dysregulated proteins are synapse-enriched protein kinases such as Src and calmodulin-dependent protein kinase IIα and IIβ (CaMKIIα and CaMKIIβ) we previously predicted to catalyse phosphorylation of specific neuronal proteins during excitotoxicity (Hoque et al., 2019). Additionally, we demonstrate in a rodent model of excitotoxicity that blocking proteolytic processing of Src can protect against excitotoxic neuronal loss. Finally, we describe how experiments to validate our TAILS findings *in vivo* in a mouse model of ischemic stroke led to the discovery of new mechanisms of regulation of CaMKIIα under both physiological conditions and during excitotoxicity. Since CaMKIIα is a drug target for protection against excitotoxic neuronal death in an acute neurological disorder (Deng, Orfila et al., 2017, Leurs, Klein et al., 2021), our discoveries suggest that blockade of proteolytic processing of dysregulated synapse-enriched protein kinases or inhibition of these dysregulated kinases could be new neuroprotective strategies to reduce brain damage in neurological disorders.

## RESULTS

### Remodelling of the neuronal N-terminome during excitotoxicity occurs without changes in protein abundance

The signaling pathways directing neuronal death upon over-stimulation of iGluRs are poorly characterized. Whilst these pathways are likely activated at an early stage following over-stimulation of iGluRs, excitotoxic cell death does not occur immediately; rather it is the prolonged activation of these pathways that causes the ultimate demise of neurons (Hoque et al., 2019, Hossain et al., 2013). One model to study these events is by treating cultured primary mouse cortical neurons with glutamate (reviewed in (Choi, 1990)). Consistent with previous reports, our experiments involving timed treatment of primary mouse neuronal cultures also demonstrated delayed cell damage and death, observed only after 240 min of glutamate treatment (Figure S1). Therefore, proteomic analysis was performed at the 30-min (early) and 240-min (late) time points after glutamate treatment to identify neuronal proteins demonstrating altered abundance or modification by proteolysis at the early and late stages of excitotoxicity. We reasoned that identifying temporal changes in neuronal proteins undergoing significantly enhanced proteolysis at these two treatment time points could unveil initiating and effector molecular events occurring during excitotoxicity. Some of these events, initiated at the early stage of excitotoxicity when neurons are still alive and sustained until late stages when neuronal death is noticeable, may represent potential drivers of excitotoxic cell death. Using the TAILS method (Kleifeld, Doucet et al., 2011), we identified and quantified over 5000 N-terminal peptides derived from neuronal proteins in all experimental conditions (Figure S2A, Tables S1A and S2A). Among them, over 70 % contain the neo-N-terminal amino acid residues generated by proteolysis of intact neuronal proteins during excitotoxicity (Figure S2A). From the identified neo-N-terminal residues and the abundance changes of the neo-N-terminal peptides, we defined the identities and mechanisms of dysregulation of specific neuronal proteins undergoing proteolytic processing during excitotoxicity.

The two major purposes of proteolysis in cells are (i) to degrade proteins into intermediate peptide fragments destined for clearance into dipeptide and amino acids (herein referred to as degradation) (Figure S2B) and (ii) to process proteins by removing regulatory domains or motifs to form stable truncated protein fragments with altered biological activities (herein referred to as proteolytic processing) (Figure S2C). Degradation of cellular proteins is mediated by proteasomal and lysosomal systems, while proteolytic processing is catalyzed by modulator proteases that specifically target cleavage sites in an intact protein to generate one or more truncated protein fragments. As most of these fragments contain intact functional domain(s), they are relatively stable and may even perform functions different from those of the intact proteins. As shown in Figure S2C, we assigned neo-N-terminal peptides whose abundance was increased as being derived from stable truncated fragments generated from enhanced proteolytic processing of neuronal proteins by excitotoxicity-activated proteases. Conversely, peptides whose abundance decreased during excitotoxicity were assigned as being derived from neuronal proteins undergoing enhanced degradation for clearance (Figure S2B). Using statistical analysis to define the thresholds for assigning the above groups of neo-N-terminal peptides (Kleifeld et al., 2011, Prudova, Gocheva et al., 2016) (Figures S3A & S3B), we found 234 and 365 peptides underwent significant changes in abundance at 30 min and 240 min of glutamate treatment, respectively, due to enhanced proteolysis of their parent neuronal proteins during excitotoxicity (Figures 1A and S3B). These peptides were further classified into those derived from proteins undergoing enhanced proteolytic processing and those derived from proteins undergoing enhanced degradation (Figure 1A; Tables S1B and S2B). These findings reveal for the first time the identities, cleavage sites, stability and consequences of proteolysis of cellular proteins targeted by excitotoxicity-activated proteases in neurons.

**Figure 1.**
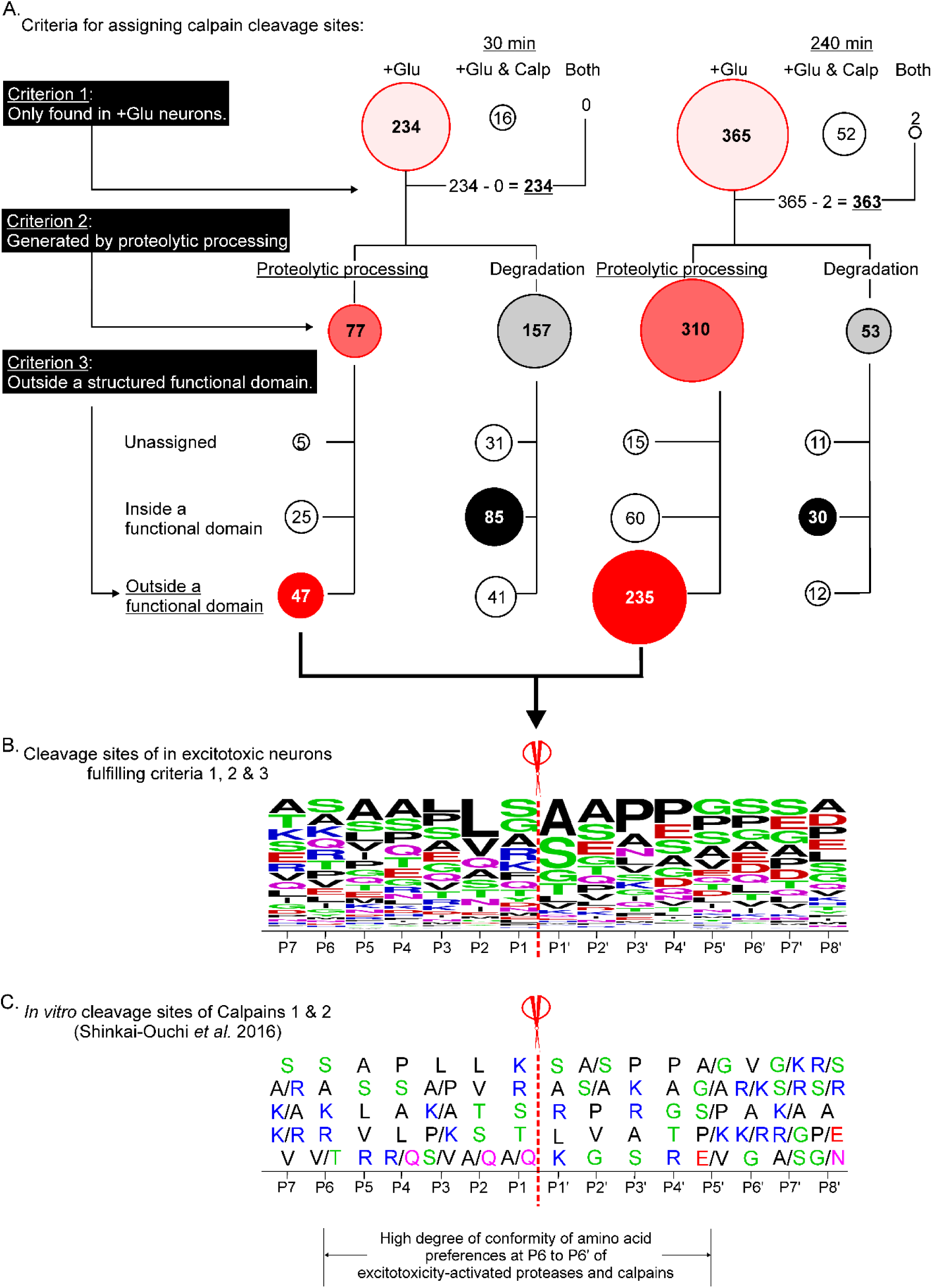
Identification of potential substrates of calpains in excitotoxic neurons. **A.** Assignment of the calpain cleavage sites from neo-N-terminal peptides identified in excitotoxic neurons. Neo-N-terminal peptides derived from neuronal proteins undergoing enhanced proteolysis induced by glutamate treatment or glutamate/calpeptin co-treatment were classified as depicted in Figures S2 and S3. The numbers of significantly changed neo-N-terminal peptides in all treatment groups assigned by sequentially applying each of the three criteria are depicted in bubble plots. The bubble size reflects the number of neo-N-terminal peptides. *Bubbles with different shades of red*: neo-N-terminal peptides found in glutamate-treated neurons only (Criterion 1), generated by proteolytic processing (Criterion 2) and predicted by Alpha fold protein structure database to reside outside a functional domain (Criterion 3). Neo-N-terminal peptides fulfilling all three criteria are assigned as those derived from neuronal proteins cleaved by calpains during excitotoxicity. *+Glu only*: peptides identified in glutamate-treated neurons; *+Glu & Calp only*: peptides identified in the glutamate/calpeptin co-treated neurons; *both*: peptides identified in both the glutamate-treated and the glutamate/calpeptin co-treated neurons. **B.** The frequencies of amino acids at positions P7 to P8’ proximal to the assigned calpain cleavage sites in excitotoxic neurons. The amino acid frequencies at each position are represented in WebLogo format (*89*). The sizes of representation of the listed amino acids in each position reflects the frequencies of its appearance in the identified sequences. **C.** The top five most frequent amino acids at each position in the P7-P8’ cleavage site sequences in synthetic peptides proteolyzed by calpain 1 and calpain 2 *in vitro* defined by Shinkai-Ouchi, *et al*. (*27*). The amino acids are presented from top to bottom in order of preference with the top-ranked most frequently encountered amino acid residue listed at the top and the fifth ranked frequently encountered residues listed at the bottom. *Bottom of panel C*: High conformity of amino acid preferences in P6-P5’ positions of the cleavage site sequences of calpains in excitotoxic neurons and those of calpains 1 and 2. Red: D and E; blue: K, R and H; black: ALVIPM; purple: Q and N; Green: S, T and G.

Of the ∼200-300 neo-N-terminal peptides derived from the significantly proteolyzed neuronal proteins during excitotoxicity, only 41 were found in neurons at both treatment time points (Figure S3B), indicating that neuronal proteins undergoing enhanced proteolysis at early stage of excitotoxicity are mostly different from those at late state of excitotoxicity. Additionally, among the 234 identified proteolyzed proteins in, only 77 were proteolytically processed to form stable protein fragments, while the 310 out of 365 of the proteolyzed proteins were proteolytically processed to form stable protein fragments (Figure 1A). Since these protein fragments with neo-N-terminus differ from their parent neuronal proteins in their regulatory properties and/or functions, our results suggest that generation of these fragments perturbs intracellular signaling in neurons at the late stage of excitotoxicity.

In contrast to the extensive changes in neuronal N-terminome revealed by TAILS analysis, data from global proteomic analysis (Figure S4) indicate that glutamate treatment for up to 240 min had little impact on neuronal protein abundance – only 1 and 13 neuronal proteins were deemed to have changed significantly in abundance at 30 and 240 min of glutamate treatment, respectively (Figure S5, Tables S3). Furthermore, none of the neuronal proteins exhibiting enhanced proteolysis during excitotoxicity showed significant changes in abundance (i.e. ≥ 2-fold changes) (Figure S6 and Table S4), supporting our prediction that the truncated neuronal fragments generated by proteolytic processing are stable (Figure S2B). Intriguingly, none of the neuronal proteins undergoing enhanced degradation during excitotoxicity showed significant decrease in abundance (Figure S6); some even exhibiting a small albeit non-significant increase in abundance. These results suggest that their enhanced degradation is likely a result of their increased turnover i.e. increased biosynthesis and degradation during excitotoxicity.

### Calpeptin abolished almost all excitotoxicity-related proteolytic changes in neuronal proteins

Since calpeptin, an inhibitor of calpains and cathepsins could protect cultured neurons against excitotoxic cell death (Wang, Zhang et al., 2015), cleavage of specific neuronal proteins by one or both of these proteases likely contributes to excitotoxic neuronal death. To identify the neuronal proteins proteolyzed by these proteases during excitotoxicity, we used the TAILS method to compare changes in the N-terminome of neurons treated with a combination of glutamate and calpeptin (+Glu and Calpeptin). Using the rationale depicted in Figure S2 to identify neo-N-terminal peptides derived from proteins undergoing significant proteolysis in the co-treated neurons (Figure S7A), we found 16 and 54 neo-N-terminal peptides derived from neuronal proteins undergoing significant proteolysis at 30 and 240 min of the co-treatment, respectively (Figures S7B & S7C). None of these neo-N-terminal peptides were derived from proteins undergoing significant proteolysis at 30 min of treatment and only two of them were derived from proteins significantly proteolyzed after 240 min of treatment in neurons treated with glutamate only (i.e. +Glu) (Figure S7C, Tables S5, S6 & S7). Hence, calpeptin treatment abolished almost all proteolytic changes that neuronal proteins undergo during excitotoxicity (Figure S7C), implicating a critical role for calpains and/or cathepsins in catalyzing these events.

### Most proteolytically processed proteins in excitotoxic neurons are potential substrates of calpains

The X-ray crystal structure of calpain-1 and the predicted structure of calpain-2 in alpha-fold protein structure database show that they both have a deep and narrow active site (Moldoveanu, Campbell et al., 2004, Tunyasuvunakool, Adler et al., 2021). To access to this active site, the cleavage site in a substrate needs to adopt a fully extended conformation and/or reside in an unstructured region located outside a functional domain (Moldoveanu et al., 2004). To identify potential substrates of calpains amongst the neuronal proteins with neo-N-terminal peptides derived from enhanced proteolysis in glutamate-treated neurons (Figure 1A), we applied three filtering criteria: (1) the neo-N-terminal peptides are present in neurons treated with glutamate but not in neurons treated with glutamate/calpeptin; (2) the neo-N-terminal peptides were derived from enhanced proteolytic processing of neuronal proteins; and (3) the neo-N-terminal peptides were derived from cleavage sites located outside a functional domain because proteolytic processing occurs in loop region or unstructured motifs connecting the properly folded functional domains (Tables S8A and S8B). Using these criteria, 47 and 235 neo-N-terminal peptides respectively, were identified as being derived from proteins potentially cleaved by calpains in neurons at 30 and 240 min of glutamate treatment (Figure 1A). Amongst these were protein kinases such as CaMKIIβ, p21-activated kinase 1, 3 and 6 (PAK1,3 and 6), and the regulatory subunit of protein kinase A (PRKARA) (Figure S8A). These findings suggest that proteolytic processing by calpains alters signaling events governed by these neuronal protein kinases. In addition, the stable truncated fragments generated by proteolytic processing contain one or more intact functional domains such as the kinase domain, suggesting that some of these fragments retain biological activities and may contribute to the neurotoxic signaling directing cell death during excitotoxicity.

Figure 1B shows the frequencies of amino acids proximal to the identified cleavage sites in potential calpain substrates selected by these filtering criteria (Figure 1A). The pattern of amino acid preferences from positions P6-P5’ of the cleavage site sequences is very similar to that of cleavage site sequences in synthetic peptides cleaved by calpain 1 and calpain 2 *in vitro* (Shinkai-Ouchi, Koyama et al., 2016) (Figure 1C). Since none of the significantly proteolytically processed proteins exhibited significant reduction in abundance (Figure S6), their cleavage by calpains during excitotoxicity likely generate stable truncated fragments (Table S8).

A protein undergoing degradation for clearance generates numerous unstructured or partially folded intermediate peptide fragments originating from both functional domains and disordered structural motifs. As such, cleavage sites of the degraded proteins do not preferentially reside in unstructured motifs outside a functional domain. Consistent with this prediction, most identified cleavage sites in significantly degraded neuronal proteins were located in functional domains with well-defined three-dimensional structures (Figure S8B and Table S8). Hence, the neo-N-terminal peptides we identified as those originating from the degraded neuronal proteins were likely derived from intermediate peptide fragments generated during the course of degradation of these proteins for clearance (Figure S8B). Calpeptin abolished proteolysis of proteins undergoing enhanced degradation in the glutamate-treated neurons (Figure S7C, Tables S5, S6 and S7) and cathepsins, which catalyze protein degradation, are activated during excitotoxicity (reviewed in (Fujikawa, 2015) and (Repnik, Stoka et al., 2012)). For these reasons, cathepsins are likely the proteases catalyzing enhanced degradation of neuronal proteins in the glutamate-treated neurons identified in our TAILS analysis.

### Proteolytic processing events during excitotoxicity are associated with perturbation of synaptic organization and function in neurons

Our results indicate that enhanced proteolytic processing in excitotoxic neurons generates stable truncated protein fragments lacking one or more functional domains of the parent protein. As such, these truncated protein fragments may possess dysregulated activities that can perturb biological processes in neurons. To define these processes, we interrogated how enhanced proteolytic processing of the identified neuronal proteins (listed in Tables S1B and S2B) affects known signaling pathways in cells using several predictive software and databases of cell signaling analysis and protein-protein interactions. First, interrogation with the ingenuity Pathway Analysis (IPA) software and database (Kramer, Green et al., 2014) revealed signaling pathways regulating synaptic processes including synaptogenesis, axonal guidance, cell junctions and cell-cell interactions as the most impacted signaling pathways in neurons during excitotoxicity (Figure S9). SynGO and X-link-MS databases documenting the locations, functions and protein complex formation of synaptic proteins (Gonzalez-Lozano, Koopmans et al., 2020, Koopmans, van Nierop et al., 2019, Szklarczyk, Gable et al., 2019) were used to predict how these proteolytically processed neuronal proteins perturb synaptic organization and function. Our analysis revealed that the proteolytically processed synaptic proteins mapped to all key synaptic components, with many forming stable protein complexes (Figures 2A and S10). More importantly, they participate in many synaptic biological processes (Figures 2A and S10). Their assignment to specific locations and biological processes of synapses as depicted in Figures 2A and S10 suggests dysregulation of the identified neuronal proteins by calpain-catalyzed proteolytic processing and their resultant aberrant signaling at synapses contributes to excitotoxic neuronal death.

**Figure 2.**
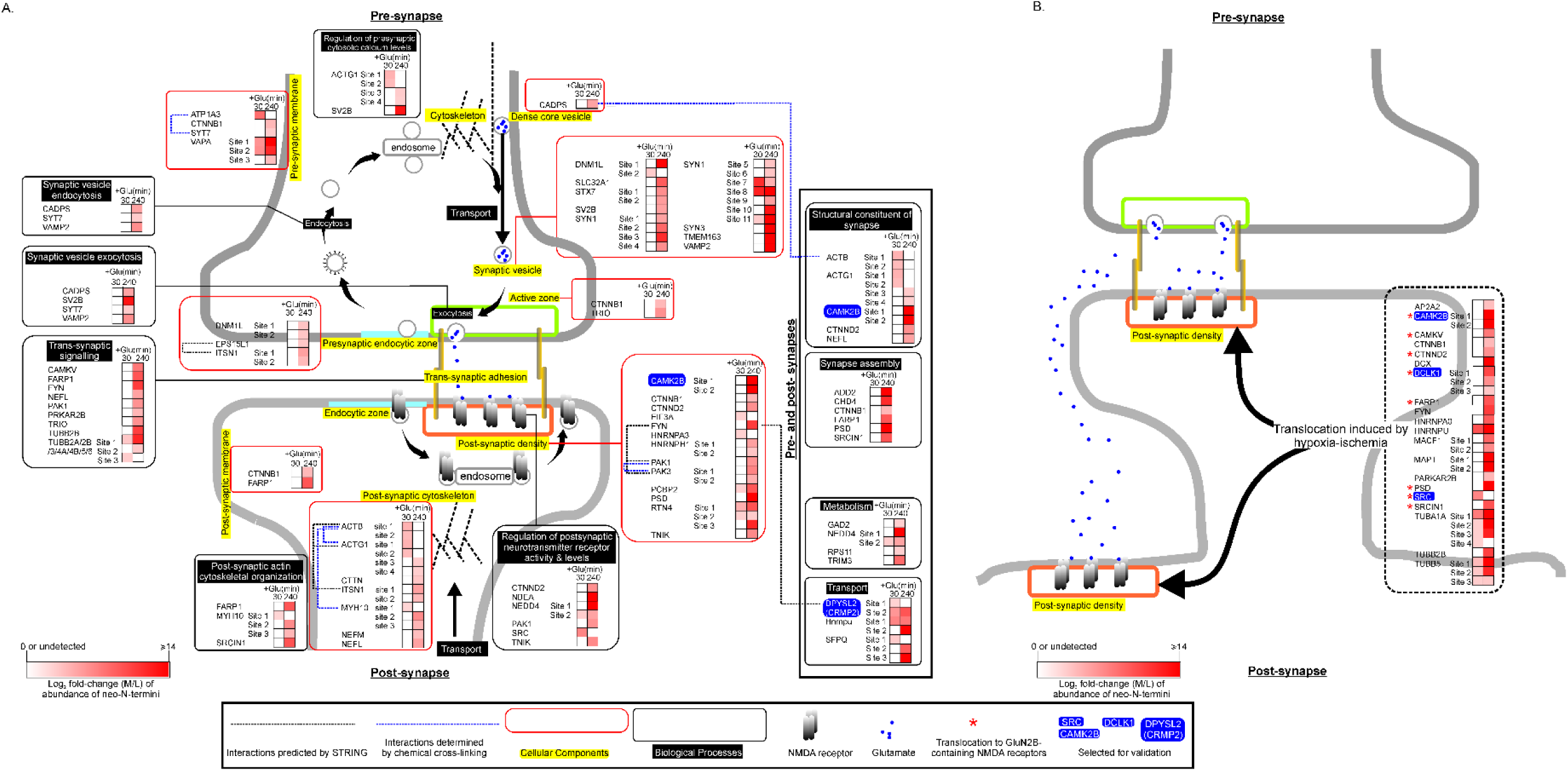
Functions and synaptic locations of proteolytically processed neuronal proteins. **A.** Neuronal proteins undergoing significantly enhanced proteolytic processing in glutamate-treated neurons were analyzed by SynGO for their synaptic locations (highlighted in yellow and grouped in red boxes) and biological processes (white fonts in black background and grouped in black boxes). *Blue dotted lines*: linking proteins that form complexes identified by cross-linking mass spectrometry by Gonzalez-Lozano, *et al*. (*30*). *Black dotted lines*: linking proteins that form complexes documented in STRING and with the complex formation confirmed experimentally. STRING parameters for analysis of the interaction networks are (i) network type: physical subnetwork, (ii) meaning of network edges: evidence, (iii) active interaction sources: experiment and database and (iv) minimum required interaction score: highest confidence (0.900). **B.** Synaptic proteins recruited to postsynaptic density in neonatal mouse brain cortex following hypoxia-ischemia (*33, 34*). Red asterisks: synaptic proteins recruited to the GluN2B-containing NMDA receptors (*33*). *Heatmap*: abundance ratios of the neo-N-terminal peptides found in glutamate-treated neurons versus those found in control neurons (presented as Log_2_-normalized M/L ratio). Neo-N-terminal peptides either undetected or showing no significant changes in abundance are depicted in white while those showing a significant increase are in red.

Twenty neuronal proteins revealed by our TAILS study to undergo enhanced proteolytic processing in excitotoxic neurons were previously found to translocate to post-synaptic density in cortex of neonatal mouse brains following hypoxia-ischemia (Lu, Shao et al., 2018, Shao, Wang et al., 2017) (Figure 2B). Among them, eight were found to be recruited to the post-synaptic density harboring GluN2B subunit-containing NMDA receptors (Lu et al., 2018) (Figure 2B). Excitotoxicity is a major mechanism directing neuronal loss in hypoxia-ischemia (Choi, 2020) and GluN2B subunit-containing NMDA receptors are NMDA receptors governing excitotoxic neuronal death (Parsons & Raymond, 2014). Based upon these findings, we postulate that these twenty neuronal proteins were recruited to the NMDA receptor-containing postsynaptic density during excitotoxicity, where they were proteolytically processed by the over-activated calpains to form stable truncated fragments. For the eight neuronal proteins previously found to be recruited to the GluN2B-containing NMDA receptors in excitotoxic neurons in mouse brain cortex during hypoxia-ischemia (Lu et al., 2018), their proteolytic processing may contribute to the cytotoxic signals emanating from NMDA receptors during excitotoxicity. As such, we selected three protein kinases (Src, DCLK1 and CAMKIIβ) amongst these 8 proteins for further biochemical analysis to illustrate the significance of our TAILS findings because their dysregulation by proteolytic processing could potentially contribute to excitotoxic neuronal death by aberrant phosphorylation of synaptic proteins. We also selected the microtubule-binding protein CRMP2, which is a known regulator of synaptic organization and functions for further biochemical analysis (Wakatsuki, Saitoh et al., 2011).

### Validation of TAILS findings I: A new mechanism of dysregulation of synaptic protein CRMP2 in excitotoxic neurons

The synaptic protein CRMP2 (also referred as DPYSL2) is a key regulator of neuronal axon guidance and synaptogenesis (Evsyukova, Plestant et al., 2013) whose dysregulation contributes to neuronal loss in neurodegenerative diseases by an unknown mechanism (Kondo, Takahashi et al., 2019, Wakatsuki et al., 2011). TAILS and Western blot analyses revealed proteolytic processing of CRMP2 to form stable truncated fragments of ∼57 kDa (Figures 3A, 3B & S11) during excitotoxicity. Specifically, CRMP2 was proteolytically processed at sites A^516^↓S^517^ and S^517^↓S^518^ in excitotoxic neurons (Figure 3A), leading to the generation of long truncated N-terminal fragments of ∼ 57 kDa and short truncated C-terminal fragments of ∼6 kDa. Furthermore, the cleavage sites lie in an unstructured region preferentially targeted by calpains (Figure 3C) (Ono, Saido et al., 2016). Accordingly, the cleavage was abolished by calpeptin (Figure S11), suggesting calpains as the upstream proteases catalyzing proteolytic processing of CRMP2 in neurons during excitotoxicity.

**Figure 3.**
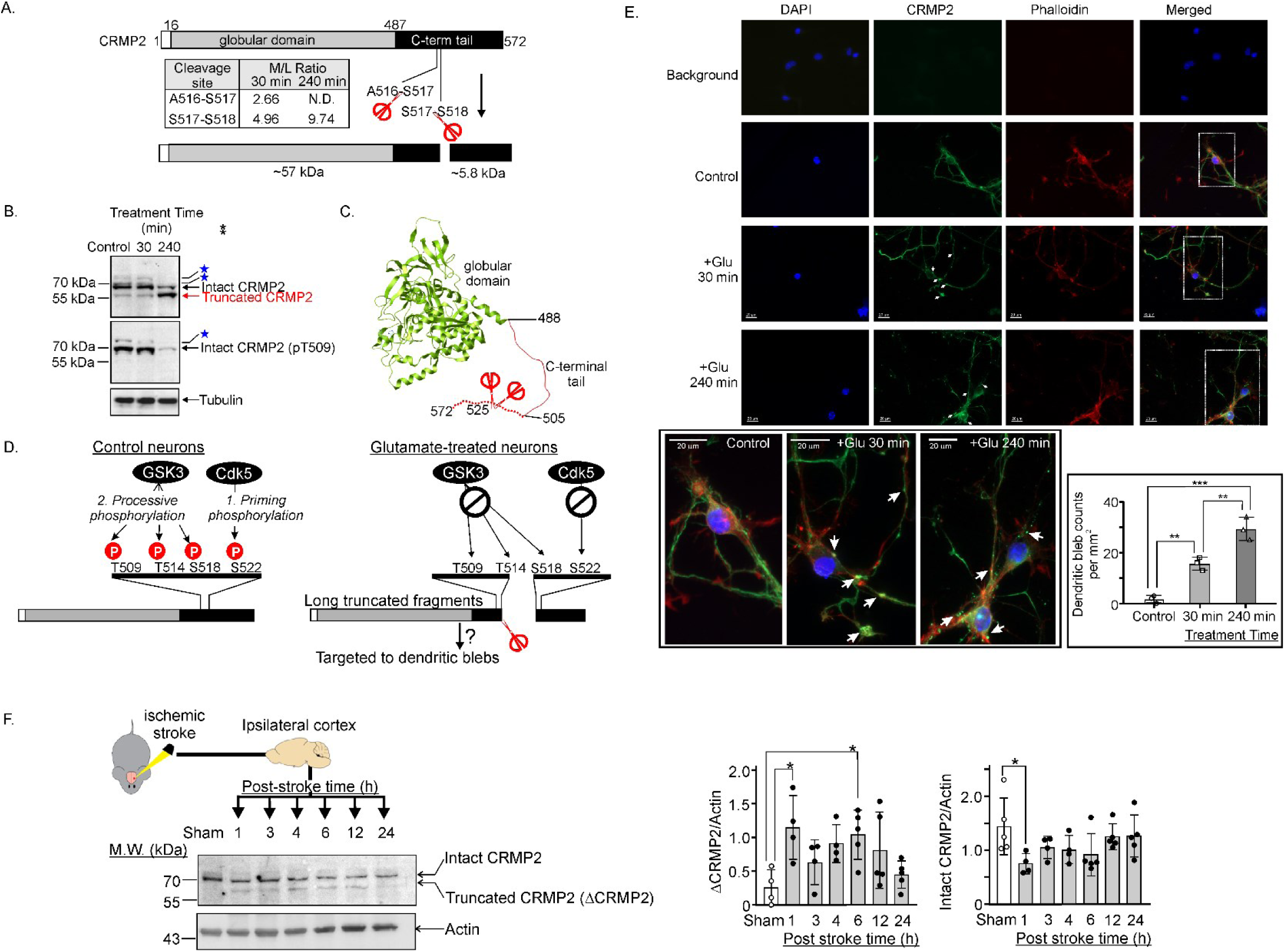
N-terminomic findings unveiled a new mechanism of dysregulation of synaptic neuronal CRMP2 (DPYSL2) **A.** CRMP2 is cleaved at sites in its C-terminal tail. Inset: the abundance (M/L) ratios of the neo-N-terminal peptides at 30 and 240 min after glutamate treatment. *N.D.*: not detected. *Red scissors*: cleavage sites. **B.** Western blots of lysates from control and glutamate-treated neurons probed with anti-CRMP2 and anti-pT509 CRMP2 antibodies. Loading control: anti-tubulin blot. *Blue stars*: potential hyper-phosphorylated forms of intact CRMP2 detected by the anti-CRMP2 and anti-pT509 CRMP2 antibodies. **C.** Structure of a phosphomimetic mutant of CRMP2 (PDB accession: 5yz5). The dotted line represents the disordered C-terminal tail region. **D.** A model depicting the new mechanism of dysregulation of neuronal CRMP2 during excitotoxicity uncovered by our findings. In control neurons, CRMP2 undergoes hierarchical phosphorylation by Cdk5 and GSK3 at sites in the C-terminal tail. Cdk5 phosphorylates the priming site S522. Upon phosphorylation, pS522 binds GSK3, which catalyzes processive phosphorylation of CRMP2 at three other sites in the order of S518, T514 and T509. In excitotoxic neurons, cleavage of CRMP2 (depicted by scissors) generates a long truncated CRMP2 fragment that lacks the priming site S522, abolishing S522 phosphorylation by Cdk5 and in turn suppressing processive phosphorylation of S518, T514 and T509 by GSK3. The truncation and lack of phosphorylation at T509, T514 and S518 may contribute to the accumulation of the immunoreactive CRMP2 signals at the dendritic blebs shown in *panel E*. **E.** Fluorescence microscopy images showing actin (phalloidin), CRMP2 and nuclei (DAPI) in control and glutamate-treated neurons. *White arrows*: dendritic blebs. *White dotted rectangles*: sections of the images selected to generate the close-up views shown in *left bottom panel*. *Right bottom panel*: number of dendritic blebs per mm^2^ in control and the glutamate treated neurons in three biological replicates. Results are presented as mean ± SD; **: p < 0.01, ***: p <0.001, one-way ANOVA with Dunnett’s multiple comparison test. **F.** Proteolytic processing of CRMP2 cortical brain tissue induced by ischemic stroke. Representative Western blot images of CRMP2 in lysates of ipsilateral brain cortex of sham operated mice and mice at the designated time points after ischemic stroke. cortical brain. The same western blot image of actin was presented in Figure 4C. The abundance ratios of intact CRMP2 and truncated (ΔCRMP2) are shown. Number of replicates: n = 4 or 5 for the sham treatment group and the ischemic stroke treatment groups at the designed post-stroke time. The blot was probed with anti-CRMP2 plus anti-rabbit Alexa 48 as primary and secondary antibodies and anti-actin plus anti-mouse Alexa 800 as the primary and secondary antibodies.

The C-terminal tail of CRMP2 contains several sites that are sequentially phosphorylated by cyclin-dependent kinase 5 (Cdk5) and glycogen synthase kinase 3β (GSK3β) in neurons (Figure 3D, left panel). In this series of events, phosphorylation of CRMP2 at S522 by Cdk5 (Uchida, Ohshima et al., 2005) directs GSK3β to subsequently phosphorylate T509, T514 and S518 in a processive manner (Uchida et al., 2005). Cleavage at A^516^↓S^517^ and S^517^↓S^518^ removes the priming phosphorylation site S522, preventing GSK3β from phosphorylating T509, T514 and S518 in the long, truncated fragment (Figure 3D, right panel). Phosphorylation of these sites impacts the ability of CRMP2 interactions with GTP-bound tubulins (Niwa, Nakamura et al., 2017, Sumi, Imasaki et al., 2018, Wilson, Ki Yeon et al., 2014), which is critical for the protein to promote axonal elongation (Inagaki, Chihara et al., 2001) and modulating microtubule dynamics (Niwa et al., 2017, Yuasa-Kawada, Suzuki et al., 2003).

Figure 3C shows that the long, truncated fragment retains the globular domain critical for promoting microtubule polymerization (Niwa et al., 2017). The microtubule polymerization-promoting activity of CRMP2 is regulated by GSK3β-mediated phosphorylation of these C-terminal tail sites (Yoshimura, Kawano et al., 2005). Owing to the lack of phosphorylation at these sites, the microtubule polymerization-promoting activity of the fragments cannot be regulated by Cdk5 and GSK3β even though both kinases are known to be aberrantly activated in neurons during excitotoxicity (Chow, Guo et al., 2014, Endo, Nito et al., 2006, Meyer et al., 2014). Consistent with our model, the truncated fragment CRMP2 fragments could not cross-react with the anti-pT509 CRMP2 antibody (Figure 3B). Collectively, the Western blot data validated our TAILS findings and prediction that CRMP2 was cleaved at sites in the C-terminal tail to generate the long-truncated N-terminal fragments that were not phosphorylated by GSK3β at T509. Based on these observations, we also predicted alteration of subcellular localization of the truncated CRMP2 fragments in excitotoxic neurons. Consistent with this prediction, glutamate treatment induced accumulation of neuronal CRMP2 in bead-like structures on dendrites called dendritic blebs (Figure 3E), previously known to form in neurons during excitotoxicity (Greenwood, Mizielinska et al., 2007). Moreover, the number of CRMP2-containing dendritic blebs in neurons at 240 min of glutamate treatment was significantly higher than that in neurons at 30 min of treatment (inset of Figure 3E). Since CRMP2 is a crucial regulator of axonal guidance signaling (Inagaki et al., 2001), our findings suggest a mechanism whereby its truncation, reduced phosphorylation at T509, and accumulation in dendritic blebs can potentially contribute to dendritic and synaptic injury associated with excitotoxic neuronal death (Hasbani, Schlief et al., 2001, Hosie, King et al., 2012).

Excitotoxic neuronal loss is a major contributor to brain damage following ischemic stroke. In agreement with our TAILS findings in excitotoxic neurons, C-terminal truncated CRMP2 fragments of ∼57 kDa were detectable at 1 – 24 h after ischemic stroke induction (Figure 3F), confirming cleavage of CRMP2 to form stable fragments *in vivo* after ischemia stroke.

### Validation of TAILS findings II: Dysregulation of synapse-enriched protein kinases by proteolytic processing during excitotoxicity

Our TAILS analysis identified several neuronal protein kinases, which were proteolytically processed during excitotoxicity to form stable truncated fragments with an intact kinase domain. Among them, Src, CaMKIIβ and DCLK1 are synapse-enriched kinases recruited to GluN2B-containing NMDA receptor in neurons in excitotoxic condition induced by hypoxia-ischemia (Lu et al., 2018) (Figure 2B). Furthermore, they play critical role in synaptic organization and/or neuronal survival (Ali & Salter, 2001, Deng et al., 2017, Hossain et al., 2013, Shin, Kashiwagi et al., 2013). As such, they were selected for further *in vitro* and *in vivo* investigations to validate our TAILS findings.

Besides its predicted role as a hub of regulation of NMDA receptor signaling in neurons (Salter & Kalia, 2004) and participation in postsynaptic organization and signaling (Figure 2), Src is also a key contributor to neuronal loss both *in vitro* and *in vivo* (Hossain et al., 2013, Liu, Waldau et al., 2017). Our TAILS study revealed for the first-time cleavage of Src at F^63^-G^64^ in glutamate treated neurons (Table S6). The abundance of the dimethyl labelled N-terminal peptide (dimethyl-G^64^-GFNSSDTVTSPQR^77^) derived from Src was 5.6-fold higher in the glutamate-treated neurons versus that in the untreated neurons (M/L ratio = 5.6) (Figure 4A, Tables S1B, S4A and S8A), indicating enhanced proteolytic processing of neuronal Src by an excitotoxicity-activated protease catalyzing cleavage of Src at the F^63^-G^64^ bond. This proteomic finding is validated by the Western blot results (Figure 4A). Second, the cleavage was predicted to generate a stable truncated fragment lacking the N-terminal myristylation domain (also referred to as the SH4 domain) and part of the unique domain (Figure 4B). This prediction was validated by the presence of a truncated fragment ΔNSrc of ∼54 kDa at 15 min till 90 min of glutamate treatment in neurons (Figure 4A) (Hossain et al., 2013) and in brain cortex of mice subjected to ischemic stroke (Figure 4C). Furthermore, recombinant Src (R-Src) cleaved by calpain 1 *in vitro* to form a 54 kDa truncated fragment as early as 2 min after incubation (Figure 4D). To determine the cleavage site in R-Src targeted by calpain 1, the reaction mixture consisting of intact R-Src only and that consisting of R-Src and calpain-1 after 120 min of incubation were subjected to isotopic dimethyl labelling prior to tryptic digestion (Figure 4D). A dimethyl labelled peptide identical to the dimethyl labelled neo-N-terminal peptide (dimethyl-G^64^-GFNSSDTVTSPQR^77^) detected in the lysate of glutamate-treated neurons, was present only in the R-Src/calpain-1 reaction mixture but not in the reaction mixture with R-Src only (Figure 4D). In summary, these results suggest that calpain 1 or another calpain isoform directly cleaves neuronal Src at the F^63^↓G^64^ bond during excitotoxicity (Figure 4B). As the resultant truncated ΔNSrc fragment lacks the regulatory myristoylation motif and unique domain (Figure 4B), its subcellular localization and kinase activity are likely dysregulated in excitotoxic neurons. In our previous study, we demonstrated that ΔNSrc mostly resided in cytosolic compartment in glutamate-treated neurons and expression a recombinant Src mutant mimicking ΔSrc in untreated primary cortical neurons led to cell death in part by inactivating the pro-survival protein kinase Akt (Hossain et al., 2013). Taken together, the results in this and our previous studies indicate that calpains directly cleave Src during excitotoxicity to generate the neurotoxic ΔNSrc, which contributes to neuronal death in part by inactivating Akt.

**Figure 4.**
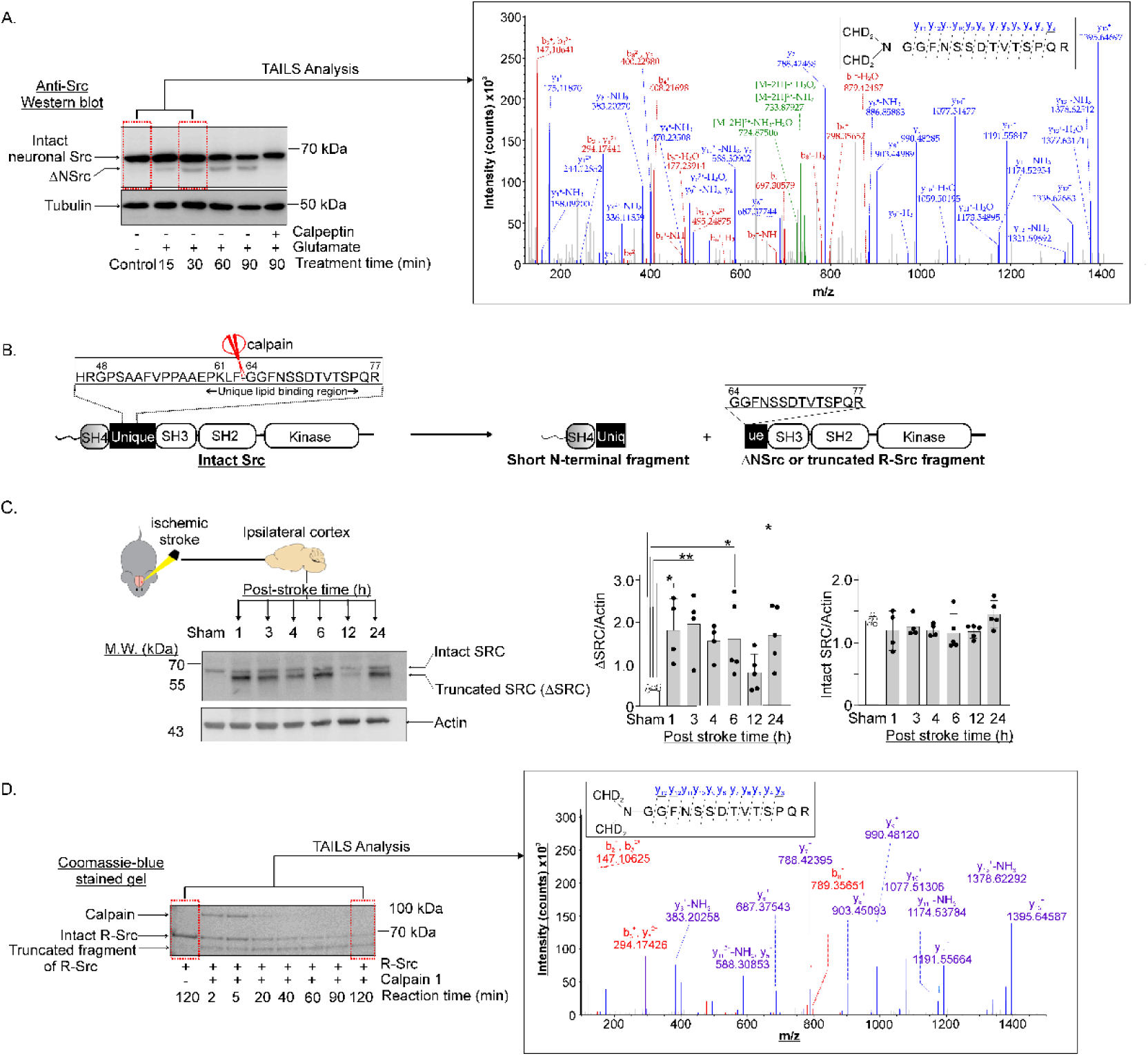
Determination of the calpain cleavage site of Src in glutamate-treated neurons and *in vitro*. **A.** Left panel: Western blot of neuronal Src in control neurons, glutamate-treated neurons and glutamate/calpeptin co-treated neurons. *ΔNSrc*: the long C-terminal fragment generated by calpain cleavage. Right panel: Fragment ion chromatogram identifying a neo-N-terminal peptide encompassing residues 54 to 77 of neuronal Src (*inset*) detectable exclusively in neurons treated with glutamate for 30 min. *Blue*: y ions, *Red*: b ions. **B.** Schematic diagram depicting the functional domains of intact Src and the calpain cleavage site (*red scissors*) and formation of a short N-terminal fragment and ΔNSrc by calpain cleavage. The cleavage site is mapped to the Unique Lipid Binding Region in the Unique domain. Hence, ΔNSrc lacks the ability to bind plasma membrane. **C.** Proteolytic processing of CRMP2 and Src in cortical brain tissue induced by ischemic stroke. Representative Western blot images of Src in lysates of ipsilateral brain cortex of sham operated mice and mice at the designated time points after ischemic stroke. The same western blot image of actin was presented in Figure 4F. The abundance ratios of intact Src and truncated Src (ΔSrc) are shown. Number of replicates: n = 4 or 5 for the sham treatment group and the ischemic stroke treatment groups at the designed post-stroke time. The same blot was probed with anti-Src antibody plus anti-mouse Alexa 800 as primary and secondary antibodies. **D.** Right panel: Coomassie blue-stained SDS-PAGE gel of reaction mixtures containing recombinant neuronal Src (R-Src) after incubation with Calpain 1 for 2 min to 120 min *in vitro*. *Boxes with dotted red lines*: samples analyzed with the TAILS method. Left panel: The fragment ion chromatogram identifying the deuterated dimethyl-labelled Src (64-77) segment of R-Src as the neo-N-terminal peptide (*inset*) detected only in the reaction mixture containing R-Src and calpain 1 at 120 min of incubation. *Blue*: y ions, *Red*: b ions.

The two CaM kinase subtypes, CaMKIIα and CaMKIIβ, are both synapse-enriched protein kinases critical to neuronal survival (Buonarati, Cook et al., 2020, Kool, Proietti Onori et al., 2019, Tullis, Buonarati et al., 2021). CaMKIIα and CaMKIIβ are highly homologous and assemble into oligomers of 12 and 14 monomeric subunits, which can consist of both subtypes (Rosenberg, Deindl et al., 2006). Although much is known about their structures, regulation, and their contribution to neuronal death during excitotoxicity (Bayer & Schulman, 2019, Buonarati, Miller et al., 2021, Guo & Chen, 2022), exactly how they direct excitotoxic neuronal death remains unclear. Our TAILS data revealed proteolytic processing of CaMKIIβ at two cleavage sites: site 1 (M^282^-H^283^) in the autoinhibitory and calmodulin binding motif and site 2 (D^389^-G^390^) in the linker motif (Figure 5A). Cleavage at sites 1 and 2 is expected to generate stable C-terminal fragments of 28.6 kDa and 17.2 kDa, respectively. Since cleavage at both sites was abolished in the glutamate/calpeptin co-treated neurons, and the cleavage sites are located in the autoinhibitory/CaM binding motif and linker motif adopting loop or poorly defined structures (Figures 5A and S8A), they are classified as direct cleavage sites of calpains by criteria depicted in Figure 1A. We next examined whether calpain-1 could cleave recombinant GST-CaMKIIβ *in vitro*. As ischemic stroke induces CaMKIIα in the mouse brain to undergo autophosphorylation at Thr-286 in the autoinhibitory/CaM binding motif (Leurs et al., 2021), we reasoned that CaMKIIβ was also autophosphorylated at the homologous site (Thr-287) in excitotoxic neurons in mouse brains subjected to ischemic stroke treatment. We therefore generated pT287 GST-CaMKIIβ by incubation of GST-CaMKIIβ with Mg^2+^-ATP in the presence of Ca^2+^/CaM and examined the cleavage product(s) of calpain-1 derived from pT287 GST-CaMKIIβ by Western blot analysis. Figure 5B shows that calpain-1 differentially cleaved the unphosphorylated and pT287-GST-CaMKIIβ and generated different C-terminal fragments. The C-terminal fragment of ∼25 kDa (C-term. ΔCaMKIIβ fragment-b in Figure 5B) derived from the unphosphorylated GST-CaMKIIβ did not correspond to any one of the two C-terminal fragments predicted from our TAILS data (Figure 5A). Intriguingly, cleavage of pT287-GST-CaMKIIβ by calpain-1 generated a fragment of ∼29 kDa (C-term. ΔCaMKIIβ fragment-a) and a fragment of ∼17 kDa (C-term. ΔCaMKIIβ fragment-c) (Figure 5B) with molecular masses corresponding to those of the C-terminal fragments generated by cleavage at sites 1 and 2, respectively as predicted by our TAILS data (Figure 5A). Figure 5C shows that ischemic stroke induced the formation of multiple C-terminal fragments of CaMKIIβ with molecular masses ranging from 26 – 35 kDa. Among them the one of ∼29 kDa and marked by a star (Figure 5C) corresponds to C-term. ΔCaMKIIβ fragment-a generated in the *in vitro* experiment (Figure 5B). Taken together, our results from these *in vitro* and *in vivo* experiments support our TAILS findings of cleavage of CaMKIIβ by calpains in neurons during excitotoxicity. Furthermore, autophosphorylation of Thr-287 facilitates calpain cleavage of CaMKIIβ at site 1 (M^282^-H^383^) mapped to the autoinhibitory/CaM binding motif (Figure 5B). Since pathological conditions leading to excitotoxic neuronal death induce autophosphorylation of CaMKII (Deng et al., 2017), these conditions are predicted to induce proteolytic processing of CaMKIIβ with site 1 being the preferred cleavage site. In agreement with this prediction, ischemic stroke induced a significant increase in the C-term ΔCaMKIIβ fragment-a.

**Figure 5.**
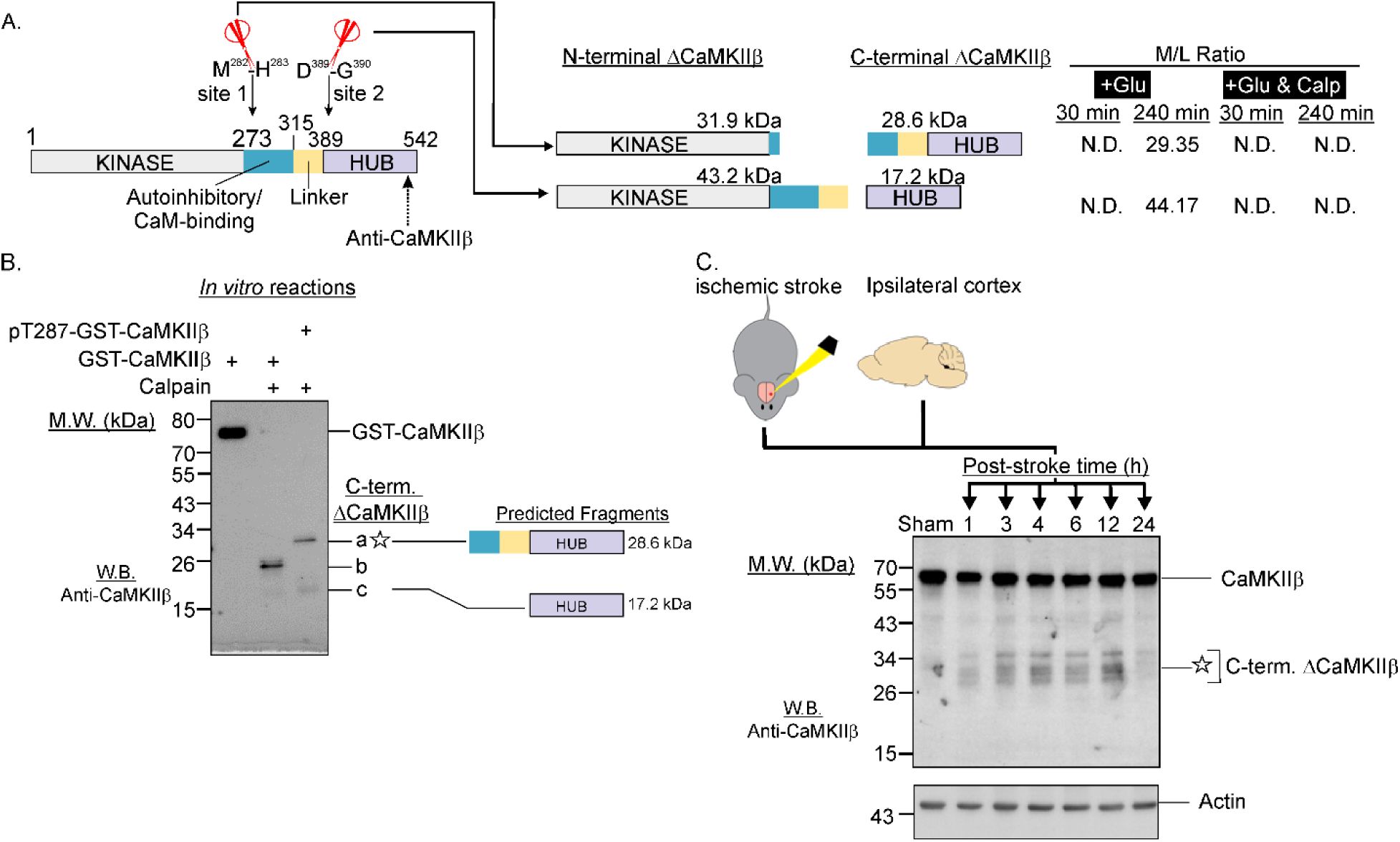
N-terminomic results led to the discovery of ischemic stroke-induced proteolytic processing of CaMKIIβ in mouse brain cortex. **A**. Schematic diagram depicting the cleavage sites in CaMKIIβ undergoing proteolytic processing in glutamate-treated neurons (+Glu) but not in neurons treated with glutamate and calpeptin (+Glu & Calp). *M/L ratio*: ratio of the neo-N-terminal peptide in the treated versus control neurons. *N.D.*: not detected. *Arrow with dotted line*: epitope of anti-CaMKIIβ antibody mapped to the C-terminal hub domain of CaMKIIβ. **B.** Differential cleavage of recombinant GST-CaMKIIβ and pT287 GST-CaMKIIβ by calpain-1. GST-CaMKIIβ with and without prior autophosphorylation at T287 (200 ng of GST-CaMKIIβ plus 100 μM ATP, 10 mM Mg^2+^, 1.5 Ca^2+^ and 5 μM CaM at 30 °C for 2 min) were incubated with calpain-1 (1 unit) *in vitro* for 45 min). The reaction mixtures were analyzed by Western blotting using the anti-CaMKIIβ antibody. *C-term. ΔCaMKIIβ*: C-terminal fragments of CaMKIIβ generated by calapin cleavage. *Predicted fragments*: Identities of C-term. ΔCaMKIIβ bands a and c predicted from N-terminomic findings shown in *panel A*. **C.** Western blot analysis of brain cortical lysates (20 μg of proteins) of sham-operated mice and mice at designated time points after ischemic stroke using the anti-CaMKIIβ antibody. Among the several C-term. ΔCaMKIIβ bands detected in ischemic stroke mouse brains, the mobility of one of them in SDS-PAGE (marked by a star) was similar to that of band a (marked by a star) in *panel B*.

DCLK1, a regulator of microtubule assembly, axonal guidance and synaptogenesis in neurons (Evsyukova et al., 2013), is a bifunctional protein consisting of two tandem doublecortin (DCX) domains and a C-terminal serine/threonine kinase domain (Gleeson, Lin et al., 1999, Reiner, Coquelle et al., 2006, Schaar, Kinoshita et al., 2004, Shu, Tseng et al., 2006). The tandem DCX domains, which drives microtubule assembly function, are connected to the C-terminal kinase domain by a large PEST sequence (linker rich in proline, glutamic acid, serine and threonine) susceptible to proteolytic cleavage (Patel, Dai et al., 2016). Calpains are known to cleave DCLK1 *in vitro* at sites mapped to the PEST motif to generate an N-terminal fragment consisting of the two DCX domains and a C-terminal fragment consisting of the intact kinase domain and a C-terminal tail (Burgess & Reiner, 2001). Consistent with these previous observations, our TAILS results revealed for the first time enhanced proteolytic processing of DCLK1 during excitotoxicity at the following sites: T^311^↓S^312^, S^312^↓S^313^ and N^315^↓G^316^ within the PEST sequence (Figure S12A). Cleavage at these sites is expected to generate N-terminal fragments of ∼35 kDa conisisting of both DCX domains and C-terminal fragments of ∼49 kDa conisisting of the kinase domain and the C-terminal tail. The cleavage was predicted to dysregulate the kinase and microtubule assembly functions of DCLK1 (Burgess & Reiner, 2001) (Patel et al., 2016) (Nawabi, Belin et al., 2015). In agreement with the TAILS findings, Western blot of the cell lysates derived from the control and glutamate-treated neurons revealed enhanced formation of truncated DCLK1 fragments of ∼50-56 kDa (Figure S12B). The epitope of the anti-DCLK1 antibody maps to the C-terminal tail of DCLK1. Hence, these truncated fragments are predicted to retain the intact protein kinase domain and the C-terminal tail. Besides detecting the 50-56 kDa truncated fragments, the antibody also cross-reacted with several truncated fragments of ∼37-45 kDa. These findings suggest that DCLK1 underwent proteolytic processing at multiple other sites in addition to the three cleavage sites identified by our TAILS analysis.

### Potential therapeutic value of our TAILS findings: blocking calpain-mediated cleavage of specific synapse-enriched proteins as a potential neuroprotective therapeutic strategy

Based on the sequence surrounding the cleavage site (F^63^-G^64^) in Src identified in our TAILS analysis (Figure 4B), we designed a cell-permeable peptide TAT-Src consisting of the cell-permeable TAT-sequence and the segment encompassing residues 49-79 in the unique domain of Src. We also designed a cell-permeable control peptide TAT-Scrambled consisting of the TAT-sequence and a segment with identical amino acid composition but scrambled sequence of the Src(49-79) region (Figure 6A). We then demonstrated that blockade of cleavage of Src by TAT-Src in excitotoxic neurons (Figure 6A). Importantly, TAT-Src but not TAT-Scrambled could protect the cultured cortical neurons against excitotoxic cell death (Hossain et al., 2013), suggesting TAT-Src as a neuroprotectant *in vitro*.

**Figure 6.**
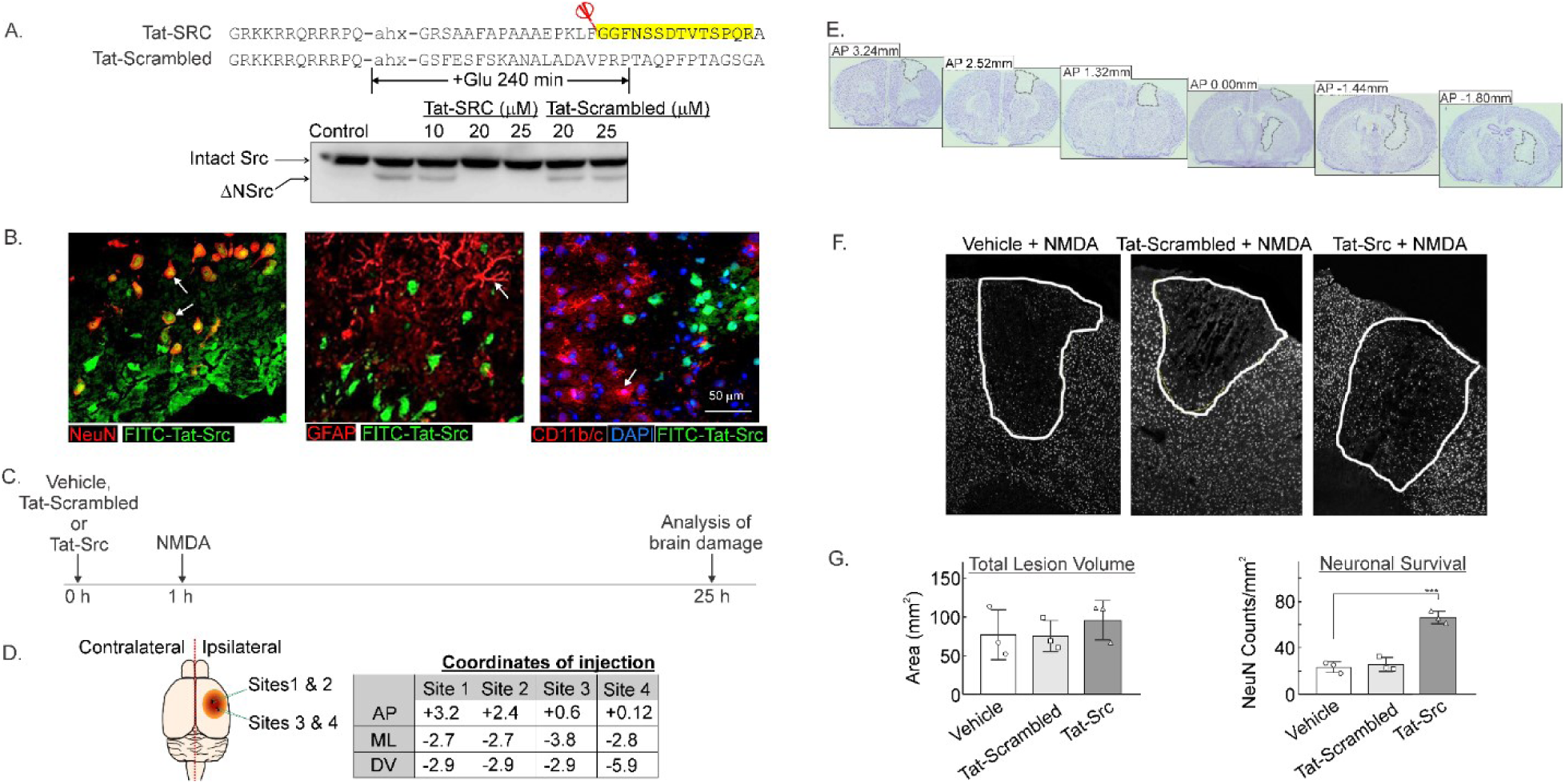
TAT-Src protects against neuronal loss *in vivo* in NMDA-mediated neurotoxicity. (**A**) Tat-SRC but not Tat-Scrambled blocks cleavage of Src in glutamate-treated neurons. *Upper panel*: Sequences of Tat-Src and Tat-scrambled-Src. Segment highlighted in yellow is the neo-N-terminal peptide corresponding to Src(64-77) detected exclusively in glutamate-treated neurons (see Figure 5A). *Red scissor*: site in neuronal Src cleaved by the excitotoxicity-activated proteases in neurons and by calpain 1 *in vitro*. *Lower panel*: Tat-Src or Tat-Scrambled of designated concentrations were added to the culture medium 1 h prior to treatment of cultured neurons with 100 μM glutamate. Cleavage of Src during excitotoxicity was monitored by anti-Src Western blot. *ΔNSrc*: truncated Src fragment. (**B**) Representative photomicrographs of FITC labelled TAT-Src infusion co-labelled against markers of neuronal cells (NeuN+ red); astrocytes (GFAP+; red) and microglia (CD11b/c OX-42+, red) and observed as orange. (**C**) Time line depicting treatment with Vehicle (Milli-Q H_2_O, 3 μl), Tat-Src (5 mM in Milli-Q H_2_O, 3 μl) or Tat-Scrambled (5 mM in Milli-Q H_2_O, 3 μl) at 1 h prior to NMDA-induced excitotoxicity. (**D**) Stereotaxic coordinates of the four injection sites (sites 1 - 4) used to cerebrally inject NMDA to induce excitotoxicity (70 mM in PBS, 1 μl per site). (**E**) Representative thionin-stained coronal images of rat brains infused with NMDA to demonstrate damage to the motor cortex and dorsal striatum. (**F**) NeuN+ cells (transposed white using Image J software) in all three treatment groups. (**G**) *Left panel*: the total lesion volumes in the treatment groups. *Right panel*: the total number NeuN+ cells in each treatment group was point-counted using image J software with the number of surviving neurons within the lesion significantly increased in rats treated with TAT-Src (*p* < 0.0001, n = 3/group, one-way ANOVA) followed by the Bonferroni post-hoc test. Data were analyzed using GraphPad Prism, version 8 and presented as mean ± SD. Statistical significance was defined as *p* < 0.05 for infarct volume and *p* < 0.0001 for NeuN cell count.

Since the TAT-Src and TAT-Scrambled peptides are peptides of 44 amino acids, it is unclear if they can pass through the blood brain barrier. We therefore used a rat model of neurotoxicity involving direct injection of NMDA and one of the two peptides to explore if TAT-Src can exert neuroprotective action *in vivo*. We first performed stereotaxic infusion of FITC-TAT-Src with a fluorescent tag covalently attached at the N-terminus of TAT-Src, into the cortical and striatal regions of rat brains (Figure 6B). FITC-TAT-Src was detected in neurons but not astrocytes and microglia in the infused regions of the rat brain (Figure 6B) one-hour post infusion, suggesting that it can enter neurons to exert its neuroprotective action. To examine the ability of TAT-Src to protect against excitotoxic neuronal death *in vivo*, TAT-Src, vehicle (water) or TAT-scrambled were stereotaxically injected at four sites in the cortical and striatal regions of rat brains (Figures 6C and 6D). One hour after the injection, neurotoxic dose of NMDA was infused to the same sites to induce excitotoxic brain damage. At 24 h after the infusion of NMDA, the rats were sacrificed, and brain sections were prepared to measure the infarct volume and the number of surviving neurons (Figure 6E). The absence, or reduction in NeuN immunoreactivity, revealed NMDA-induced lesions within the motor and parietal cortex (Figures 6E and 6F). The plot in left panel of Figure 6G shows that total lesion volume was consistent across all treatment groups with no significant difference in the volume of damage detected between groups. Stereological point counting of NeuN positive cells within the lesion revealed treatment-specific effects where the number of neurons in rats treated with TAT-Src was significantly higher than in rats receiving vehicle or scrambled Tat-Src control (Figure 6G, right panel). Thus, injection of TAT-Src prior to NMDA infusion could protect against excitotoxic neuronal loss caused by the injected NMDA. Since FITC-TAT-Src entered neurons but not astrocytes and microglia (Figure 6B), the ability of TAT-Src to protect against neuronal loss in NMDA-induced brain damage is presumably attributed to its blockade of calpain cleavage of neuronal Src to form the neurotoxic truncated Src fragment (ΔNSrc) (Hossain et al., 2013). In conclusion, our results illustrate blockade of pathologically activated proteolytic events as a neuroprotective strategy to reduce brain damage in neurological disorders.

### Extrapolation of our TAILS findings led to the discovery of new regulatory mechanisms of CaMKIIα in normal and pathological conditions

The *CaMK2b* transcript encoding CaMKIIβ was detectable in embryonic and neonatal mouse brains while *CaMK2a* encoding CaMKIIα was undetectable in these tissues (Bayer, Lohler et al., 1999). As such, CaMKIIβ expression is expected to be much higher than CaMKIIα expression in the cultured differentiated embryonic cortical neurons used in our study, explaining why TAILS analysis of the differential embryonic cortical neurons detected the neo-N-terminal peptides derived from CaMKIIβ but not those from CaMKIIα (Table S2). Given the high degree of sequence similarity of CaMKIIα and CaMKIIβ (Figure 7A), we predicted that CaMKIIα in adult neurons is also directly cleaved by calpains at the homologous sites 1 and 2 (Figure 7A) to generate two kinase domain-containing N-terminal fragments: CaMKIIα-short and CaMKIIα-long encompassing residues 1 to 282 and 1 to 389 of CaMKIIα, respectively. To validate our prediction, we incubated calpain-1 with unphosphorylated recombinant CaMKIIα (rCaMKIIα) and pT286-rCaMKIIα *in vitro*. Figure 7B shows differential proteolytic processing of unphosphorylated rCaMKIIα and pT286-rCaMKIIα. A fragment with molecular mass (∼36 kDa) similar to that of ΔCaMKIIα-long was generated by calpain-1 cleavage of both forms of ΔCaMKIIα, while a fragment with molecular mass (∼31 kDa) similar to that of ΔCaMKIIα-short was generated when pT286-rCaMKIIα was cleaved by calpain-1 (Figure 7B). These results suggest that calpain-1 targeted site 2 of both autophosphorylated and unphosphorylated rCaMKIIα. However, site 1 of rCaMKIIα was accessible to calpain-1 only when Thr-286 was autophosphorylated.

**Figure 7:**
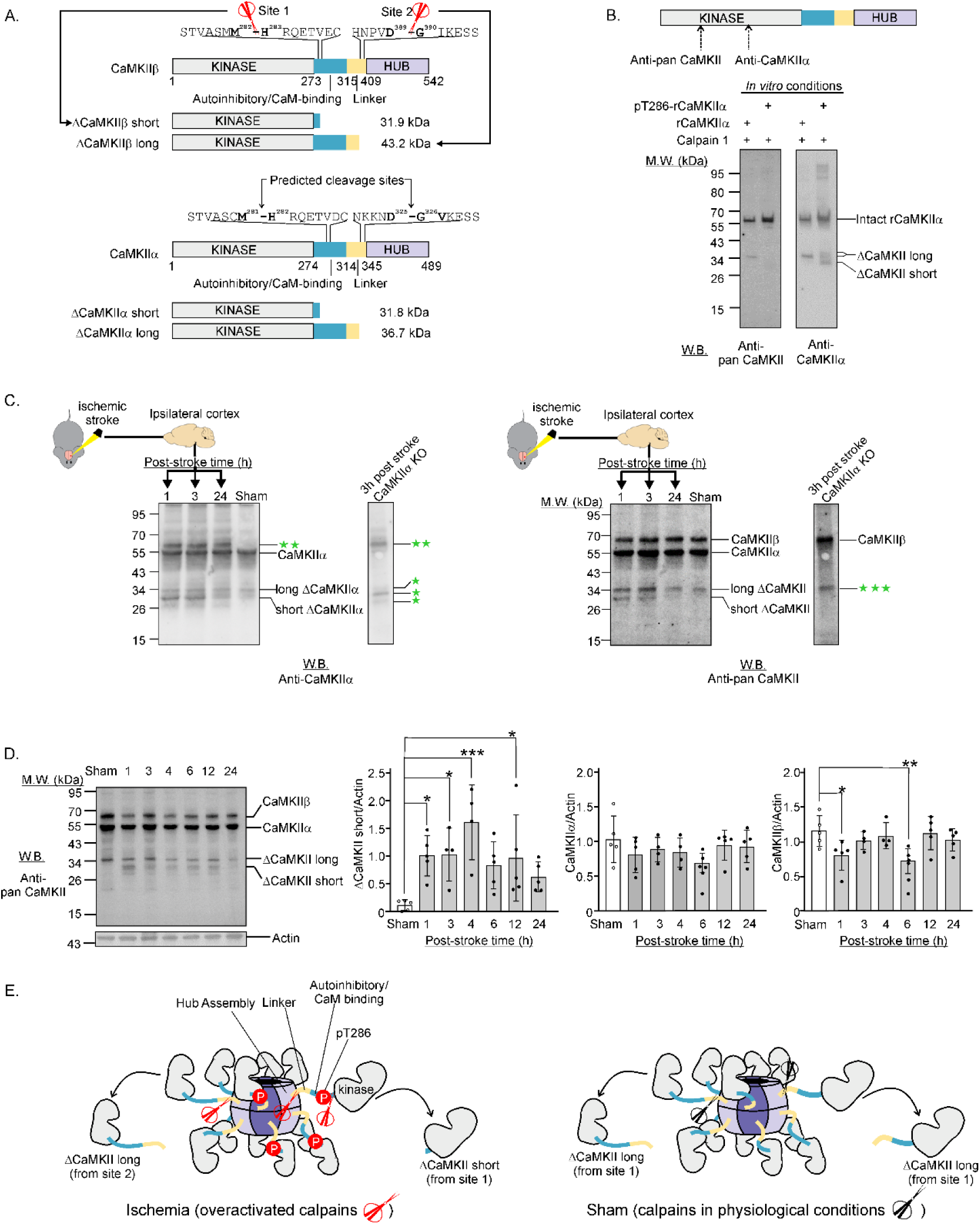
N-terminomic results contributed to the discovery of ischemic stroke-induced proteolytic processing of CaMKIIα in mouse brain cortex. **A.** Calpain cleavage sites of CaMKIIβ in excitotoxic neurons identified by TAILS and the corresponding predicted cleavage sites in CaMKIIα. *ΔCaMKII short and ΔCaMKII long*: N-terminal truncated fragments derived from cleavage at site 1 and site 2, respectively. **B.** Cleavage of unphosphorylated recombinant CaMKIIα (rCaMKIIα) and pT286-rCaMKIIα by calpain-1 *in vitro* generated truncated CaMKIIα (ΔCaMKIIα) fragments with molecular masses similar to those in ischemic stroke mouse brains. The pT286-rCaMKIIα was generated by incubating 200 μg of rCaMKIIα for 2 min at 30 °C in the presence of 10 μM ATP, 10 mM Mg^2+^, 1.5 mM Ca^2+^ and 5 μM CaM. For the *in vitro* cleavage experiments, rCaMKIIα or pT286-rCaMKIIα was incubated with calpain-1 (1 unit) for 45 min. Reaction mixtures of the *in vitro* experiments were probed with the anti-Pan CaMKII and anti-CaMKIIα antibodies with epitopes mapped to the kinase domain. **C.** Western blots of lysates of brain cortex (20 μg of proteins) collected from wild type sham-operated mice and wild type mice and CaMKIIα KO mice at designated time points after photothrombotic stroke. The blots were probed with anti-CaMKIIα antibody (*left panel*) and anti-pan CaMKII antibody. One green star: Proteins in CaMKIIα KO brain lysates recognized by the anti-CaMKIIα antibody. These proteins were likely the intact and N-terminal proteolytic fragments of other CaMKII isoforms. Two green stars: A protein band cross-reacted with anti-CaMKIIα antibody but not with anti-pan CaMKII antibody. Three green stars: the molecular mass (∼35 kDa) of this long N-terminal fragment (ΔCaMKII long) differs from the predicted molecular mass (43.2 kDa) of ΔCaMKII long derived from CaMKIIβ. As this fragment was presented in the CaMKIIα KO brain lysate, it is likely derived from another CaMKII isoform such as CaMKIIδ expressed in brain cells. **D.** Western blot of lysates of brain cortex collected from wild type sham-operated mice and at designated time points after photothrombotic stroke treatment. The blot was probed with anti-pan CaMKII antibody (*left panel*). The abundance ratios of the intact CaMKIIα or CaMKIIβ versus actin and those of the short N terminal fragment of CaMKII (ΔCaMKII short) in brain cortical lysates of sham-operated and ischemic stroke mice (*right panels*). The *p*-values are represented as * for *p*-value < 0.05, ** for *p*-value < 0.01 and *** for *p*-value < 0.001. **E**. A model of differential cleavage of CaMKII oligomer by calpains in sham and ischemic stroke mouse brains. Under physiological conditions, calpains (black scissors) cleave CaMKIIα and CaMKIIβ homo- and hetero-oligomers at site 2 to generate ΔCaMKII long containing the kinase domain, the autoinhibitory/CaM binding motif and the linker motif. In ischemic stroke condition, calpains are over-activated and the CaMKII oligomers undergo autophosphorylation at Thr-286. The over-activated calpains (red scissors) cleave the autophosphorylated CaMKIIα and CaMKIIβ oligomers at sites 1 and 2, respectively to generate ΔCaMKII short and ΔCaMKII long fragments.

Similar to the fragments generated by calpain-1 cleavage of pT286-rCaMKIIα *in vitro*, N-terminal fragments of CaMKIIα with molecular masses similar to ΔCaMKIIα-short (∼31 kDa) and ΔCaMKIIα-long (∼36 kDa) were detectable in brain cortex of mice subjected to ischemic stroke treatment (Figure 7C, left panel). Ischemic stroke induced an increase in the abundance of ΔCaMKIIα-short but not ΔCaMKIIα-long, suggesting enhanced cleavage of CaMKIIα at site 1 (M^201^-H^202^) induced by ischemic stroke. Intriguingly, the ΔCaMKIIα-long exists in brain cortex of both sham-operated mice and mice subjected to ischemic stroke treatment, suggesting that proteolytic processing at site 2 (D^925^-G^926^) of CaMKIIα occurred in both physiological and pathological conditions.

To examine if other neuronal CaMKII isoforms including CaMKIIδ and CaMKIIγ were also proteolytically processes by calpains to generate the long and short N-terminal truncated fragments (ΔCaMKII-long and ΔCaMKII-short), we probed the wild type and CaMKIIα knock-out cortical brain lysates with a pan-CaMKII antibody known to cross-react with all CaMKII isoforms (Figure 7C, right panel and Figure 7D). Since ΔCaMKII-short was detectable in brain cortex of wild type mice but not in CaMKIIα KO mice subjected to ischemic stroke treatment (Figure 7C, right panel), CaMKII-short in wild type ischemic stroke mouse brains were likely mainly derived from cleavage of CaMKIIα at site 1 by calpains.

In summary, these results demonstrate cleavage of CaMKIIα by calpains at site 2 in both physiological and ischemic stroke conditions to generate ΔCaMKIIα-long with intact kinase domain and the autoinhibitory/CaM-binding motif. Upon autophosphorylation at Thr-286 induced *in vivo* by ischemic stroke and *in vitro* by incubation with Mg^2+^-ATP and CaM, site 1 of CaMKIIα was cleaved by calpains to generate ΔCaMKIIα-short consisting of just the intact kinase domain and a remnant segment of the autoinhibitory/CaM-binding motif. Since CaMKIIα is critical to excitotoxic neuronal death (Ashpole, Song et al., 2012, Rostas, Hoffman et al., 2017), ΔCaMKII-short generated in glutamate-treated neurons is a potential neurotoxic mediator of excitotoxic neuronal death.

## DISCUSSION

We have for the first time identified over three hundred neuronal proteins associated with synaptic organization and functions that undergo enhanced proteolytic processing to generate stable truncated fragments during excitotoxicity. Since most truncated fragments derived from these proteolytically processed proteins consist of one or more intact functional domains, they likely retain some biological activities of the intact parent neuronal proteins, while other domains are perturbed. Presumably, interplay between these dysregulated truncated fragments alters synaptic organization and functions, which in turn directs neurons to undergo excitotoxic cell death. Hence, these discoveries form a conceptual framework for future investigation to define specific protein-protein interaction networks contributing to neuronal death in excitotoxicity. For example, our findings provided a shortlist of specific pharmacological tools including inhibitors of the proteolytically processed protein kinases and substrate-specific calpain inhibitors such as the neuroprotective TAT-Src peptide (Figure 6) for future investigation to chart signaling pathways orchestrating excitotoxic neuronal death.

One approach to decipher these pathways is through combinatorial phosphoproteomic and N-terminomic analysis of glutamate-treated neurons with and without co-treatment with specific inhibitors of pathologically activated cellular events occurring during excitotoxicity. These include: (i) nerinetide, a cell permeable peptide interfering over-activation of nNOS (Aarts, Liu et al., 2002) and NOX2 (Chen, Brennan-Minnella et al., 2015) by the hyperactivated NMDA receptor (Cook, Teves et al., 2012, Sun, Doucette et al., 2008), (ii) specific inhibitors of the synapse-enriched protein kinases including Src, CaMKIIα, CaMKIIβ, DCLK1, Fyn, PAK1 and PAK3 dysregulated by proteolytic processing during excitotoxic neurons (listed in Table S8), (iii) specific inhibitors of calpains 1 and 2 (reviewed in (Baudry & Bi, 2016)) and (iv) substrate-specific inhibitors of calpains such as TAT-Src (Figure 6).

We previously demonstrated that ΔNSrc generated by calpain cleavage of Src is a mediator of the neurotoxic signals of the over-stimulated NMDA receptors. Since ΔNSrc is a protein tyrosine kinase, identifying the proteins directly phosphorylated by ΔNSrc in neurons is an appropriate approach to define its neurotoxic mechanism. The proteomic method developed by Bian, *et al*. to define the phosphotyrosine-proteome of cultured cells using the superbinder mutants of SH2 domains (Bian, Li et al., 2016) is therefore a method of choice to identify the substrates of ΔNSrc in neurons during excitotoxicity.

### Sequences in proteins proteolytically processed in excitotoxic neurons contain structural features directing their cleavage by calpains

The mechanism of substrate recognition by calpains in cells are poorly understood because of the relative scarcity of confirmed calpain cleavage site data. Since the cleavage sites in the neuronal proteins identified by our TAILS analysis were the exact sites targeted by calpains in neurons, they therefore represent genuine cleavage sites of calpain substrates *in vivo*. Our data demonstrates that in excitotoxic neurons, the pattern of amino acid preferences from positions P6-P5’of the cleavage site sequences in neuronal proteins proteolytically processed by calpains is very similar to that in synthetic peptides cleaved by calpain 1 and calpain 2 *in vitro* (Figure 1). These findings suggest that the active site and substrate-binding sites (exosites) of calpains recognize specific structural features in the P6-P5’ amino acid residues of the cleavage site sequences in their *in vivo* substrates.

duVerle and Mamitsuka documented around 100 experimentally confirmed calpain cleavage sequences in mammalian proteins (duVerle & Mamitsuka, 2019). Among them, the exact sites of calpain cleavage sites were determined by *in vitro* studies. As such, it remains unclear if the cleavage sites they determined correspond to the sites of cleavage in these proteins by calpains *in vivo*. Based upon the limited data of calpain cleavage sites in protein substrates and results of peptide library studies to define the optimal cleavage sequences (Cuerrier, Moldoveanu et al., 2005, Shinkai-Ouchi et al., 2016), a number of algorithms for prediction of calpain substrates and the cleavage sites were designed. The most notable algorithms include calCleaveMKL (duVerle & Mamitsuka, 2019), iProt-Sub (Song, Wang et al., 2019), DeepCalpain (Liu, Yu et al., 2019) and GPS-CCD (Liu, Cao et al., 2011). Here, we demonstrated for the first time a high degree of conformity of amino acid preferences at the P6-P5’ positions in both the potential calpain substrates in neurons and the *in vitro* peptide substrates of calpains 1 and 2, which are the major calpain isoforms expressed in neurons (Figures 1B and 1C). Of the 200-300 cleavage sites identified in our study (Figure 1), over 90% were identified for the first time as potential calpain cleavage sites in properly folded proteins in live cells. As such, incorporating information of these newly identified cleavage site sequences and/or three-dimensional structures of the corresponding calpain substrates will improve the predictive accuracy of these algorithms.

Besides the primary structure proximal to the cleavage site and the three-dimensional structural features, co-localization of a potential calpain substrate with a specific isoform of calpains also governs whether it is proteolytically processed *in vivo*. For example, the C-terminal tail of calpains 1 and 2 contains different PDZ-binding motifs which target them to different subcellular compartments where they proteolyze specific subsets of protein substrates (Baudry & Bi, 2016, Wang, Briz et al., 2013). Future investigations to decipher where and when the potential calpain substrates identified in our TAILS analysis form protein complexes with calpains 1 and 2 in neurons during excitotoxicity will further bridge the knowledge gap concerning how the two isoforms of calpains recognize their substrates in neurons.

### Neuronal proteins proteolytically processed during excitotoxicity are potential targets for the development of neuroprotective therapeutics

Currently, there is no FDA-approved neuroprotective drug for treating patients suffering from acute neurological disorders or neurodegenerative disease. This pessimistic scenario has been challenged by the promising results of two clinical trials of a cell membrane permeable peptide nerinetide, which inhibits a key pathological event directing excitotoxic neuronal death – binding of the scaffolding protein PSD95 to the NMDA receptor (Aarts et al., 2002, Hill et al., 2020, Hill, Martin et al., 2012). The positive clinical outcomes of nerinetide treatment illustrates that other events occurring in neurons during excitotoxicity are potential therapeutic targets for the development of neuroprotective strategies for the treatment of ischemic stroke patients. Proteolytic processing of Src to generate the neurotoxic truncated fragment ΔNSrc is also a pathological event directing neuronal death (Hossain et al., 2013) and we demonstrated the substrate-specific calpain inhibitor Tat-Src could protect against neuronal cell loss *in vivo* (Figure 6), small-molecule compounds mimicking TAT-Src peptide in specifically blocking calpain cleavage of Src in neurons (Figure 6) are potential neuroprotective drug candidates to reduce excitotoxic neuronal loss in neurological disorders.

It is noteworthy that not all substrate-specific cell membrane-permeable peptides blocking neuronal proteins from being proteolytically processed by calpains are potential neuroprotectants. For example, cell-permeable peptide inhibitors that specifically block calpain cleavage of CaMKIIα and CaMKIIβ at sites 1 and 2 (Figure 7) are not suitable for use as neuroprotective therapeutics. Cleavage site 1 (M^282^-H^283^ of CaMKIIβ) is located within the autoinhibitory/CaM-binding motif. Hence, even if a cell-permeable peptide derived from the sequence around this cleavage site can specifically block calpain cleavage of CaMKIIβ in excitotoxic neurons, it would not be suitable for use as a neuroprotectant, as the peptide sequence contains the autoinhibitory/CaM-binding motif, which can bind CaM and interfere with the physiological function of CaM *in vivo*. Thus, careful peptide sequence optimization to eliminate its CaM-binding ability would be needed. Since autophosphorylation of CaMKIIα at Thr-286 impairs its CaM-binding activity, replacing Thr-286 in the peptide sequence with a phosphomimetic amino acid may achieve this aim. For site 2 (D^389^-G^390^ of CaMKIIβ), proteolytic processing of CaMKIIβ and CaMKIIα at this site occurs in both sham-operated and ischemic stroke mouse brains (Figure 7), indicating that it is a proteolytic processing event occurring in both physiological and pathological conditions. Hence, the cell-permeable peptide derived from the sequence around this site is not suitable for use as a neuroprotective therapeutic because it can potentially interfere with modifications of CaMKIIα and CaMKIIβ by calpains under physiological conditions. It is thus necessary to define the effects of blockade of calpain cleavage of the proteolytically processed neuronal proteins identified in our TAILS analysis before they are chosen as the targets for future development of neuroprotective therapeutics. Nonetheless, besides identifying calpain cleavage of Src as a potential target for development of neuroprotective therapeutics, the proteolytic processing events during excitotoxicity constructed by our TAILS analysis also provide the conceptual framework for identifying other calpain substrates as potential targets for neuroprotective therapeutics development.

### Calpain-catalyzed proteolytic processing as a physiological and pathological regulatory mechanism of neuronal CaMKII

Results of Western blot analysis indicated that cleavage of CaMKIIα at site 2, leading to the formation of ΔCaMKIIα-long consisting of the intact kinase domain and the autoinhibitory/CaM-binding motif, occurred in both sham-operated and ischemic stroke mouse brains (Figures 7C, 7D and 7E), suggesting proteolytic processing as a physiological regulatory mechanism of CaMKII. In contrast, ΔCaMKII-short, consisting of just the intact kinase domain was generated by cleavage of CaMKIIα and other neuronal CaMKII isoforms at site 1 was generated in ischemic stroke mouse brains only (Figures 7C, 7D and 7E). Hence, proteolytic processing of CaMKIIα and other isoforms at site 1 leading to the formation of ΔCaMKII-short is a pathological regulatory mechanism of CaMKII.

Autophosphorylation at Thr-286 of CaMKIIα and the homologous threonine residue in other CaMKII isoforms allows CaMKII to remain active without stimulation by Ca^2+^/CaM (Bhattacharyya, Karandur et al., 2020, Miller & Kennedy, 1986). As Thr-286 and the homologous threonine residues in CaMKII isoforms are located in the autoinhibitory/CaM-binding motif, suggesting that its phosphorylation can affect the accessibility of both cleavage sites 1 and 2 to calpains. Results of *in vitro* analysis revealed that cleavage at both sites occurred only when Thr-286 and Thr-287 of CaMKIIα and CaMKIIβ, respectively were autophosphorylated (Figures 5B and 7B). This effect of phosphorylation at Thr-286/Thr-287 on calpain-catalyzed proteolysis of CaMKIIα and CaMKIIβ was in agreement with results of the *in vitro* studies reported by Rich, *et al*. (Rich, Schworer et al., 1990) and Kwiatkowski and King (Kwiatkowski & King, 1989). These findings suggest that autophosphorylation at Thr-286/Thr-287 is a pre-requisite for calpain-mediated cleavage of CaMKII at both sites 1 and 2.

Besides enhanced proteolytic processing by calpain cleavage at site 1, CaMKIIα and its isoforms including CaMKIIβ and CaMKIIδ expressed in neurons (referred to as CaMKIIs) undergo significant autophosphorylation at Thr-286. Upon phosphorylation of Thr-286 and its homologous threonine, CaMKIIs exhibit autonomous Ca^2+^/CaM-independent activity. How might autophosphorylation at Thr-286 and calpain cleavage at site 2 contribute to excitotoxic neuronal death and brain damage in ischemic stroke? Using the mouse model of cardiac arrest and pulmonary resuscitation (CA/CPR) that cause excitotoxic neuronal loss, CA/CPR induced significant brain damage and a significant increase in autophosphorylation at Thr-286 and the homologous threonine of CaMKII and other isoforms in synaptosome membrane fractions (Deng et al., 2017). Treatment with the tight binding CaMKII inhibitor tatCN19o at 30 min after CA/PCR reduced autonomous CaMKII activity and reduced the brain damage inflicted by the treatment, suggesting that the neuroprotective action of tatCN19o is attributed to its inhibition of autonomous kinase activity and/or autophosphorylation at Thr-286 of CaMKII. Further investigation with the transgenic mice expressing the phosphorylation-deficient T286A-CaMKII mutant revealed that, similar to the effects of tatCN19o, CA/CPR inflicted much less brain damage. In summary, these findings confirm the neurotoxic role of sustained Ca^2+^/CaM-independent activation of CaMKIIs as a result of autophosphorylation at Thr-286 and the homologous threonine residues. In light of our TAILS and biochemical findings (Figures 5 and 7), we hypothesize that CaMKIIs autophosphorylated at Thr-286 and homologous threonine residues in excitotoxic neurons are cleaved by calpains at site 1, leading to the formation of the constitutively active ΔCaMKII-short, which phosphorylates specific neuronal proteins to direct neuronal death (see model in Figure 7E). Future investigation into the effects of recombinant ΔCaMKII-short on neuronal survival will reveal whether ΔCaMKII-short is a mediator of excitotoxic neuronal death. Of relevance, we previously demonstrated that the truncated fragment ΔNSrc generated by proteolytic processing of Src was a mediator of excitotoxic neuronal death (Hossain et al., 2013). Additionally, a recombinant truncated CaMKII fragment with an intact kinase domain was highly active and its expression in neurons induces AMPA receptor activation in synapses (Hayashi, Shi et al., 2000), confirming the ability of truncated fragment of CaMKII to induce aberrant signaling events in neurons.

What are the physiological consequences of calpain cleavage of CaMKIIs at site 2 to generate ΔCaMKII-long? Since the CaMKII-long fragments consist of the intact kinase domain and the autoinhibitory/CaM-binding motif, their activity is still under the control of Ca^2+^/CaM. However, lack of the hub domain allows CaMKII-long to dissociate from the CaMKII oligomeric complexes, translocate to a different subcellular location and phosphorylate specific neuronal proteins in that location. Future investigations to identify the substrates and binding partners of CaMKII-long fragments in neurons will shed light on their potential physiological functions.

## MATERIALS AND METHODS

### Experimental Design

#### Animals

Both mice (C57BL/6, C57BL/6J and *Camk2a^-/-^*) and rats (male, male Hooded Wistar) were used in both the *in vitro* and *in vivo* studies. For preparation of cultured mouse cortical neurons, the procedures were approved by the University of Melbourne Animal Ethics Committee (License number: 161394.1) and were performed in accordance with the Prevention of Cruelty to Animals Act 1986 under the guidelines of the National Health and Medical Research Council Code of Practice for the Care and Use of Animals for Experimental Purposes in Australia.

For global proteomic and N-terminomic analyses of mouse cortical neurons, cultured mouse neurons derived from mouse embryos were used. The embryos were collected from pregnant C57BL/6 mice (gestational day 14-15) after they were euthanized by CO_2_ asphyxiation.

For the dual carotid artery ligation (DCAL) ischemic stroke mouse model and the controlled cortical impact mouse model of traumatic brain injury (TBI), male C57BL/6 mice (20-30 g) were used. Brains from sham-operated mice and those from mice subjected to dual carotid artery ligation ischemic stroke and traumatic brain injury were prepared solely for the construction of spectral libraries. All experiments were performed in strict accordance with the guidelines of the National Health & Medical Research Council of Australia Code of Practice for the Care and use of Animals for Experimental Purposes in Australia.

For the animals used in the mouse photothrombotic stroke models, young male C57BL/6*J* (3-4 months, 27-30 g) were obtained from the Biomedical Research Facility, University of Otago, New Zealand. *Camk2a^-/-^*mice (*Camk2*^atm3Sva^, MGI:2389262 mice backcrossed into the C57BL/6*J* background) were bred in the *In Vivo* Pharmacology Research Unit, University of Copenhagen, Denmark. All procedures on wildtype C57BL/6*J* mice were performed in accordance with the guidelines on the care and use of laboratory animals set out by the University of Otago, Animal Research Committee and the Guide for Care and Use of Laboratory Animals (NIH Publication No. 85–23, 1996). For stroke surgeries involving *Camk2a^-/-^* mice ethical permission was granted by the Danish Animal Experiments Inspectorate (permission 2017-15-0201-01248), and all animal procedures were performed in compliance with Directive 2010/63/EU of the European Parliament and of the Council, and with Danish Law and Order regulating animal experiments (LBK no. 253, 08/03/2013 and BEK no. 88, 30/01/2013). All procedures were performed in accordance with the ARRIVE (Animal Research: Reporting In Vivo Experiments) guidelines. All measures were taken to minimize pain, including subcutaneously administration of buprenorphine hydrochloride (0.1 mL of a 0.5 mg/kg solution, Temgesic®) as pre-emptive post-surgical pain relief.

For the animals used in the rat model of neurotoxicity, male Hooded Wistar rats weighing 300-340g sourced from Laboratory Animal Services, University of Adelaide, Australia were used. The protocol was approved by the St Vincent’s Hospital animal ethics committee (AEC016/12). All surgery was performed under general anesthesia, and paracetamol (2 mg/kg in drinking water) was provided for 24 h prior to and after surgery in order to minimize suffering and included monitoring each rat throughout the length of the study to ensure their wellbeing.

#### Experimental Model I - Preparation of cultured mouse primary cortical neurons

Cultured mouse cortical neurons were prepared for the construction of spectral libraries, multi-dimensional proteomic analyses and validation of the proteomic results by biochemical methods. The cortical region was aseptically micro-dissected out of the brains of the embryos, free of meninges and digested with 0.025 % (w/v) trypsin in Krebs Buffer (0.126 M NaCl, 2.5 mM KCl, 25 mM NaHCO_3_, 1.2 mM NaH_2_PO_4_, 1.2 mM MgCl_2_, 2.5 mM CaCl_2_, pH 7.2) and incubated at 37°C with shaking to dissociate the connective tissues. After 20 min of incubation, 0.008% (w/v) DNase I (Roche Applied Science) and 0.026 % (w/v) soybean trypsin inhibitor (Sigma) in 10 ml Krebs solution (DNase I/soybean trypsin inhibitor solution) were added to the suspension to stop the trypsin action and initiate digestion of DNA. Gentle mixing by inversion of the suspension was performed. Cells in the suspension were collected by centrifugation at 1000 × g for 3 min at room temperature. They were resuspended in 1 ml of DNase I/soybean trypsin inhibitor solution. After aspiration of the supernatant, the cells were resuspended in plating medium (minimum essential medium (MEM) supplemented with 2 mM L-glutamine, 0.22% v/v NaHCO3, 1% penicillin-streptomycin, 5% v/v horse serum and 10% v/v fetal calf serum). Approximately 800,000 cells per well were seeded to a 12-well plate pre-treated with 0.1 mg/ml Poly-D-lysine. After incubation at 37°C in 5% CO_2_ for 2 h, the medium was replaced with neurobasal medium supplemented with 0.2 mM L-glutamine, 0.01 mg/ml penicillin-streptomycin and B27 supplement (NB/B27). Cultures were maintained for seven days (days *in vitro* 7 (DIV7)). Immunofluorescence microscopy analysis using antibodies against neuronal, astrocytic and microglial markers revealed that the DIV7 culture contained 94.1 ± 1.1% neurons, 4.9 ± 1.1% astrocytes, <1% microglia and <1% other cells (data not shown). The DIV7 cultures, highly enriched with neurons, were used for the experiments.

To induce excitotoxicity, the DIV 7 neuronal cultures were treated with 100 μM glutamate in NB/B27 for 30 and 240 min. For co-treatment with glutamate and calpeptin, the DIV 7 neuronal cultures were treated with 100 μM glutamate and 20 μM calpeptin in NB/B27 for 30 and 240 min. For control, viable untreated cells from the same batch of DIV 7 neuronal cultures were used for proteomic and biochemical analyses.

#### Experimental Model 2 - Mouse models of DCAL ischemic stroke and traumatic brain injury used for proteomic analyses

To generate the DCAL stroke model, mice were placed into a plastic box and initially anaesthetized with 5% Isoflurane (in 1.0 ml/min O_2_), and maintained on 1.5% Isoflurane for the duration of the experiment. A temperature probe connected to a thermoblanket (Harvard Apparatus Ltd., Kent, UK) was inserted into the rectum to monitor body temperature, and body temperature was maintained at 37°C throughout the experiment using a heat lamp. All surgical procedures were performed using aseptic technique. Surgical instruments were sterilized using a bead sterilizer (Steri350, Sigma Aldrich) prior to use. The mouse was affixed in a supine position on the thermoblanket using tape, and the neck shaved and swabbed with alcohol. Through a ventral midline incision, the carotid arteries were exposed via blunt dissection and carefully dissected clear of the vagus nerve and surrounding tissue. A stabilization period of 10 min is allowed between the isolation of each artery. The right jugular vein was also exposed via a cut to the skin and blunt dissection for the purpose of drug administration. Following the surgical procedures, the animals were allowed to stabilize for 10 min before the experiment proceeded. The left carotid artery is permanently ligated using 6-0 silk suture (6-0 black braided silk suture, SDR Scientific, Sydney, Australia). Following a stabilization period of 10 min, the right carotid artery is then transiently occluded for 30 min using a small hemostatic clamp. After the transient occlusion, the neck incision was closed using tissue glue (Leukosan Adhesive, BSN medical Inc.), and the mouse allowed to recover under 1% O_2_. Animals were housed separately following surgery with access to food and water *ad libitum*. At 24 h post-surgery, the mice were euthanized, and their brains removed and stored at -80 ⁰C.

A mouse model of traumatic brain injury (TBI) was used to generate brain lysates to construct the spectral libraries for global proteomic analysis. For this model, controlled cortical impact (CCI) was induced. This procedure is well established and induces a reproducible brain trauma with a mortality of less than 5% (Sashindranath, Samson et al., 2011). After anesthesia with 0.5 g/kg avertin (1.875% solution dissolved in 0.9% sodium chloride pre-warmed at 37°C, 2,2,2-tribromoethanol; Sigma Aldrich #T48402; 1mg/ml in tert-amyl alcohol), injected intra-peritoneally (i.p.), mice were placed in a stereotaxic frame (Kopf, Tujunga, CA). A sagittal incision in the midline of the head was performed and the skull cleaned with a 6% hydrogen peroxide solution using a sterile cotton swab. This was followed by a 5 mm diameter craniotomy performed with a drill over the left parietal cortex. The impactor was positioned in a 20° angle with the tip (cylindrical rod) touching the brain surface, and a single blunt force trauma was inflicted to the exposed brain area with an impact depth of 2mm, a velocity of 5 m/s and dwell time of 400 ms inducing a moderate to severe brain trauma. The exposed brain was then sealed using bone wax (Ethicon, Johnson and Johnson #W810T) and the skin incision was sutured with a non-absorbable braided treated silk (Dynek, Dysilk) and treated with a local anesthetic (xylocaine) and an antiseptic (betadine). For the sham procedure, only the scalp incision, without craniotomy and CCI was performed since even the craniotomy without CCI results in a brain lesion (Sashindranath, Daglas et al., 2015). Regardless of the experimental design, however, mice were placed on a 37 °C heat pad within 30 min after induction of anesthesia for post-surgery recovery, and they usually recovered within 40-60 min. Animals were housed separately following surgery with access to food and water *ad libitum*. At 24 h post-surgery, the mice were euthanized, and their brains removed and stored at -80 ⁰C.

To prepare the brain lysates from DCAL ischemic stroke mice, TBI mice and sham-operated mice for generation of the spectral libraries, we homogenized the frozen tissue biopsies in ice-cold RIPA buffer (50 mM Tris-HCl, pH 7.0, 1 mM EDTA, 5 mM EGTA, 1 mM dithiothreotol, 10% (v/v) glycerol, 1 % Triton X-100, 0.01% SDS, 150 mM NaCl, 50 mM NaF, 40 mM sodium pyrophosphate, 0.5 mM Na_3_VO_4_, 50 mM β-glycerol phosphate, 0.2 mM benzamidine, 0.1 mg/ml phenylmethyl sulfonyl fluoride (PMSF)) supplemented with 1% cOmpleteTM protease inhibitor cocktail (Roche Diagnostics). The tissue lysates were harvested and centrifuged at 12500×g for 15 min at 4 °C. Supernatants were collected and the protein concentrations were determined by BCA protein assay (Pierce-Thermo Scientific) prior to storage at -80 °C for further processing.

#### Experimental Model 3 – A mouse model of photothrombotic ischemic stroke used for biochemical investigation of proteolytic processing of CaMKII, Src and CRMP2

For the mouse model of photothrombotic stroke used for biochemical investigation of proteolytic processing of CaMKII, Src and CRMP2, young adult male C57BL/6J mice (3-4 months, 27-30 g, *n* = 4-5) were subjected to photothrombotic stroke as previously described (Clarkson, Huang et al., 2010). In brief, mice were anesthetized with isoflurane (4% induction, 2-2.5% maintenance in O2) and body temperature was kept at 37 °C using a heating pad throughout the procedure. Mice were placed in a stereotactic frame (9000RR-B-U, KOPF; CA, USA). The skin was sterilized using chlorhexidine (30% in 70% ethanol, Hibitane®). Following exposure of the skull through a midline incision, it was cleared of connective tissue and dried. A cold light source (KL1500 LCD, Zeiss, Auckland, New Zealand) attached to a 40x objective providing a 2- mm diameter illumination was positioned 1.5 mm lateral from bregma. Next, 0.2 mL of Rose Bengal (Sigma-Aldrich, Auckland, New Zealand; 10 mg/mL in normal saline) was administered i.p 5 min prior to illumination. Then, the brain was illuminated through the exposed intact skull for 15 minutes with 3300 K color temperature intensity. The skin was glued and animal were returned to their home a cage placed on a heating pad during the wake-up phase. Sham surgery was performed in the exact same way, except that saline was injected instead of Rose Bengal. Mice were housed in groups of two to five under standard conditions in individually ventilated cages (IVC: Tecniplast): 21 ± 2 °C and humidity of 50 ± 10%, on a 12 h light/dark cycle with ad libitum access to food and water. Mice were monitored and weighed after surgery. All animals were randomly assigned to experimental groups. No deaths were reported during these studies. One mouse was excluded from stroke + 4 h survival group due to the lack of any visible stroke being detected, most likely due to experimenter error with the Rose Bengal most likely being injected into the bladder. Mice were euthanized by cervical dislocation, followed by rapid extraction of the brain at 1, 3, 4, 6, 12 and 24 h after stroke induction. Brains were snap-frozen and stored at -80 °C until further processing.

For preparation of the brain lysate from photothrombotic ischemic stroke mice, brains were processed and prepared for Western blot analysis as previously described in Leurs *et al* (Leurs et al., 2021). In brief, peri-infarct tissue was collected at - 20 °C using a tissue punch, and tissue homogenization was performed using a Bullet Blender in RIPA buffer supplemented with 1% cOmpleteTM protease inhibitor cocktail (Roche Diagnostics), 1% phosphatase inhibitor cocktail 2 (#P5726, Sigma-Aldrich) and 1% phosphatase inhibitor cocktail 3 (#P0044, Sigma-Aldrich). Protein concentration was determined with the Pierce™ BCA Protein Assay Kit (#23227, Thermo Fisher Scientific). Samples were prepared for Western blot analysis by addition of 4× Fluorescent Compatible Sample Buffer (#LC2570, Thermo Fisher) and 100 mM DTT with a protein concentration of 2 μg/μL. Samples were heated for 5 min at 95 °C, sonicated and centrifuged 11,000 x g for 2 min at 4 °C. 20 μg sample were loaded onto 4-20% Mini-PROTEAN® TGXTM gels (Bio-Rad), and SDS-PAGE was performed for 40 min (200 V) with 1x Tris/glycine/SDS (25 mM Tris, 192 mM glycine, 0.1% SDS, pH 8.3) running buffer. Protein transfer to a PVDF membrane (#4561096, Biorad) was performed using the Trans-Blot® TurboTM transfer system (Bio-Rad) (2.5A, 7min), and membranes were blocked 1x BlockerTM FL Fluorescent Blocking Buffer (#37565, Thermo Fisher) for 30 min at room temperature.

Following primary antibody incubation, membranes were washed 3x 5 min washes in 1x tris-buffered saline (TBS) with 0.1% Tween-20 detergent (TBS-T). Next, membranes were probed with species-specific secondary antibodies, and washed again 3x for 10 min with TBS-T. Image were aquired with the iBright FL1500 imaging system (Invitrogen), and signals were quantified in Image Studio (Lite version 5.2). Data was analyzed in GraphPad Prism (version 8), presented as mean ± SD, and statistical analysis was performed using One-way ANOVA, post-hoc Dunnett’s test.

#### Experimental Model 4 - in vivo model of NMDA neurotoxicity

To induce NMDA excitotoxicity *in vivo*, male hooded Wistar rats (n = 12) weighing 300-340g (Laboratory Animal Services, University of Adelaide, Australia) were used. Rats were anesthetized with intraperitoneal administration of ketamine and xylazine (75mg/kg and 10 mg/kg, respectively), and maintained by inhalation of isoflurane (95% oxygen and 5% isoflurane). Rats were positioned in a stereotaxic frame (Kopf), and 4 burr holes were drilled into the right hemisphere corresponding to the predetermined sites for NMDA infusion (Site 1: AP +3.2, ML - 2.7, DV -2.9; Site 2: AP +2.4, ML -2.7, DV -2.9; Site 3: AP +0.6, ML -3.8, DV -2.9; Site 4: AP +0.12, ML -2.8, DV -5.9). Rats were randomly assigned into 3 cohorts and received either vehicle (sterile Milli-Q H_2_O, 3 μl per site), TAT-SRC peptide (5 mM in sterile MilliQ H_2_O, 3 μl per site), or TAT-Scrambled peptide (5 mM in sterile MilliQ H_2_O, 3 μl per site) via direct infusion at 0.2 μl/min into each site 1 h prior to infusion of NMDA (70 mM, 1μl PBS per site). Following infusion each burr hole was filled with bone wax and wounds sutured. In separate studies, rats were infused with FITC-TAT-SRC peptide without NMDA to assess neuronal uptake and cell specificity at one hour after infusion.

For analysis of the effect of TAT-SRC on neuronal loss in vivo, randomization was used in group allocation and data analysis. Rats that received treatment with NMDA +/- TAT-SRC, TAT-Scrambled or Vehicle were allowed to recover for 24 h prior to lethal overdose (lethobarb) and decapitation. Forebrains were collected and rapidly frozen over liquid nitrogen and stored at -80°C prior to processing. Coronal sections (16 µm thick) were prepared using a Leica cryostat (Leica Microsystems, Wetzlar, Germany) across the 4 coronal planes corresponding to the NMDA infusion sites.

For immunohistochemistry analysis, immunofluorescence staining was performed in forebrain tissue sections to identify TAT-SRC cell specificity and treatment effects following NMDA excitotoxicity. Sections were first fixed in 4% PFA for 15 minutes at RT prior to wash (3 × 5 min washes with 0.1 M PBS) and a blocking step in 5 % NGS and 0.3 % Triton X-100 and 0.1 M PBS for 30 min. Sections were again washed (3 × 5 min washes with 0.1 M PBS) and adjacent sections incubated with primary antibodies to detect neurons, astrocytes and microglia using the following primary antibodies: mouse anti-NeuN (1:500, Chemicon); mouse anti-GFAP (1:400, Millipore); and mouse anti-OX-42 (1:100, Serotec) in 2% NGS, 0.3% Triton X-100 and 0.1M PBS overnight at 4°C. The following day sections were again washed (3 × 5 min washes with 0.1 M PBS) and incubated with secondary fluorophore-conjugated antibody Alexa 555 goat anti-mouse (1:500, Thermo Fisher Scientific) for visualization of each primary antibody. For all experiments, DNA counterstain DAPI (Molecular Probes, Thermo Fisher Scientific) was applied before coverslipping with ProLong gold anti-fade reagent (Invitrogen). Control studies included omission of each primary antibody.

For lesion assessment and stereology of rat brains, triplicate sections from each NMDA infusion site were visualized using an Olympus microscope (Albertslund, Denmark) equipped with a 578–603 nm filter set for the detection of red fluorescence, at ×20 magnification. NMDA-induced lesions were identified by a distinct reduction or absence of NeuN fluorescence, which was analyzed manually by tracing the site of injury using ImageJ software (NIH, Bethesda, MD, USA). Lesion volume was then determined as described by Osborne, *et al*. by integrating the cross-sectional area of damage between each stereotaxic infusion site and the distance between sites (Osborne, Shigeno et al., 1987). The number of NeuN positive cells within each lesion were also point counted using ImageJ software using a grid overlay to estimate the total number of NeuN positive cells within each region, and expressed as number of cells/mm^2^. Data obtained for infarct volume and the effects of treatment on neuronal counts were analyzed by one-way ANOVA followed by the Bonferroni post-hoc test. For infarct volume analysis, based on an a priori power analysis for one-way ANOVA (G*Power 3.1.9.2), we used at least 3 animals per group to find a large effect size on reduced size, where 40 % reduction was considered improved with the standard deviation of 20% (f = 0.6, α = 0.05, power = 0.80). For stereology to quantify the number of surviving neurons, based on a priori power analysis for one-way ANOVA (G*Power 3.1.9.2), we used at least 3 animals per group to find a large effect size on surviving neurons, where 20% salvage is considered improved and the standard deviation is 8 % (f = 0.2, α = 0.05, power = 0.80). Data were analyzed using GraphPad Prism, version 6 (GraphPad Software Inc., San Diego) and presented as mean ± standard error of the mean (SEM). Statistical significance was defined as *p* <0.05.

For investigation of cell specificity of TAT-SRC peptide in rat brains, Immunofluorescence within adjacent tissue sections was visualized using a fluorescent microscope equipped with a 578–603 nm filter set for detection of red fluorescence (NeuN, GFAP, OX-42), a 495–519 nm filter set for the detection of green fluorescence (FITC-TAT-SRC), and a 478-495nm filter set for detection of blue fluorescence (DAPI) (ZeissAxioskop2, North Ryde, Australia). Immunohistochemical co-localization of the stereotaxically infused FITC-TAT-SRC was clearly identified in the cortex and striatum and was co-localized with the neuronal marker NeuN but not with the astrocytic marker GFAP or the microglial marker OX-42 one-hour post-infusion.

For infarct assessment, absence or reduction in NeuN immunoreactivity in brain sections was monitored because it revealed NMDA-induced lesions within the motor and parietal cortex as well as the striatum. Total lesion volume was consistent across treatment groups with no significant difference in volume of damage detected between groups (P >0.05, n=3/group, one-way ANOVA). Stereological point counting of NeuN positive cells within the lesion revealed treatment specific effects where the number of neurons in rats treated with TAT-SRC was significantly greater than in rats receiving vehicle or TAT-Scrambled control (P<0.0001, n=3/group, one-way ANOVA).

#### MTT cell viability assay and LDH release assay of neuronal death

Primary cortical neurons were incubated for 480 min with and without the addition of 100 μM of glutamate. The control neurons were incubated for 480 min in culture medium. For neurons treated with glutamate for 30 min, 60 min, 120 min and 240 min, they were pre-incubated in culture medium for 450 min, 420 min, 360 min and 240 min, respectively prior to the addition of glutamate to induce excitotoxicity. For neurons treated with glutamate for 480 min, they were treated with glutamate without pre-incubation in culture medium.

Cell viability was determined from primary cortical neurons (seeded in 24-well plates) using the 3-(4,5-dimethylthiazole-2-yl)-2,5-diphenyltetrazolium bromide (MTT) assay. MTT stock solution (5 mg/ml (w/v) in sterile PBS) was diluted 1/100 in culture medium. At the end of treatment of cultured neurons, the culture medium was aspirated and replaced by the diluted MTT solution. After incubation for 2 h, the diluted MTT solution was removed and 200 µl DMSO was added per well to dissolve the formazan crystals formed. Absorbance at 570 nm was measured using the Clariostar Monochromator Microplate Reader (BMG Lab Technologies, Durham, NC). Cell viability was expressed as a percentage of the control cells.

The activity of LDH released from the damaged neurons to the culture medium was measured. Briefly, 50 μl of culture medium from each well of the culture plate was transferred to 96 well-microtiter plates (Falcon). 100 μl of LDH assay mixture containing equal volume of LDH assay substrate solution, LDH Assay dye solution and LDH assay cofactor was then added to each well. The reaction was allowed to proceed at room temperature for 30 min in the dark and was stopped by adding 50 μl of 1 mM acetic acid. The absorbance at 490 nm of whole mixture was measured in triplicate the Clariostar Monochromator Microplate Reader (BMG Lab Technologies, Durham, NC). The release of LDH was calculated as a percentage of the untreated control.

#### Fluorescence histochemical analysis of cultured cortical neurons

For fluorescence histochemistry of cultured primary neurons, isolated cells were seeded onto poly-lysine treated 12 mm glass coverslips (placed in 12 well plates) and allowed to mature for 7 days in culture before glutamate treatment. Following treatment, cells were fixed in 4% paraformaldehyde for 20 min, permeabilized (10% goat serum in PBS containing 0.01% Triton-X) for 20 min and then blocked (10% goat serum in PBS) for 60 min. Cells were incubated with anti-CRMP2 antibody (diluted 1:1000 in block buffer) overnight at 4°C, then PBS washed, and incubated in an anti-rabbit-Alexa488 secondary antibody (1:500 in block buffer), anti-phalloidin-Tritc (1 µM, Sigma) and DAPI (1 µM, Sigma) for 60 min and a final PBS wash before being mounted onto glass slides using antifade mounting media (Prolong Gold, Invitrogen). A Zeiss axioscope2 microscope using a 40X objective lens equipped with Zeiss HRc camera with filter sets for FITC (green) and Rhodamine (red) were used to take images for histological analysis. Identical settings and exposure time were used to capture images for both peptides. Images were processed using Zen blue software (Zeiss, Germany) to generate tiff images before importing them into ImageJ software (NIH, version 1.5b) and the number of blebs were counted in fields.

#### Cleavage of recombinant CaMKII by calpain-1 in vitro

For *in vitro* cleavage experiments of CaMKII, recombinant CaMKII was autophosphorylated prior to cleavage. Experiments were performed in digestion buffer (50 mM Tris-HCl, pH 7.4, 2 mM DTT, 30 mM NaCl, 10 mM CaCl_2_). To generate pT286-CaMKIIα and pT287-GST-CaMKIIβ, 200 ng of the respective purified protein was stimulated with 10 μM ATP, 10 mM Mg^2+^, 1.5 Ca^2+^ and 5 μM CaM at 30°C for 2 min. CaMKII proteins with and without prior autophosphorylation were incubated with calpain-1 (1 unit) for 45 min at 30 °C in digestion buffer, and digested sample were immediately analyzed by Western blot analysis as descried above.

#### Construction of spectral libraries of cultured cortical neurons and mouse brain tissues

For a data-dependent acquisition (DIA) based mass spectrometry experiment, it is essential we generate a library of identifiable peptides first using a standard data-dependent acquisition (DDA) approach (Figure S4). For the DIA type experiment to work, the identified peptides have to be in that library first. Excitotoxicity is a major mechanism of neuronal loss caused by ischemic stroke and traumatic brain injury. We therefore included the brains of sham-operated mice, brains of mice suffering DCAL ischemic stroke and traumatic brain injury to construct the spectral libraries and that is why the library contains pooled samples from the representative samples. Pre-fractionation of the pooled peptides was also performed to increase the number of identifiable peptides and generate a deeper library. Once we generated that library, all samples are analysed individually as a separate DIA experiment. The DIA approach then makes use of the generated library for identification and quantitation. This methodology allows for deeper identification and lower number of missing values.

Prior to quantitative analysis on the mass spectrometer, we built a spectral library, which contains all distinctive information of each identified peptides such as its retention time, charge state and the fragment ion information. The success of the quantitation largely depends on the peptides to be quantified being present in this spectral library (Ludwig, Gillet et al., 2018). As such, we pooled lysates from neurons with and without glutamate (100 µM) treatment for 30 min and 240 min together with brain lysates of mouse stroke and traumatic brain injury models. Briefly, proteins in the lysates were precipitated with cold acetone (-20°C) and then resuspended in 8 M urea in 50 mM triethylammonium bicarbonate (TEAB). Proteins are then reduced with 10 mM tris(2-carboxyethyl) phosphine Hydrochloride (TCEP), alkylated with 55 mM iodoacetamide, and digested with trypsin (trypsin to protein ratio of 1:50 (w/w)) overnight at 37°C. The resultant tryptic peptides were purified by solid phase extraction (SPE) (Oasis HBL cartridge, Waters). For global proteome analysis, 100µg of these peptides were fractionated into 8 fractions using the high pH reversed-phase fractionation kit (Pierce) according to the manufacturer’s protocol before analysis on the Q-Exactive Orbitrap.

The LC system coupled to the Q-Eaxctive Orbitrap mass spectrometer was equipped with an Acclaim Pepmap nano-trap column (Dinoex-C18, 100 Å, 75 µm x 2 cm) and an Acclaim Pepmap RSLC analytical column (Dinoex-C18, 100 Å, 75 µm × 50 cm). After pre-fractionation with the high pH reversed-phase fractionation kit, tryptic peptides in each of the 8 fractions were injected to the enrichment column at an isocratic flow of 5 µl/min of 2 % (v/v) CH_3_CN containing 0.1 % (v/v) formic acid for 6 min before the enrichment column was switched in-line with the analytical column. The eluents were 0.1 % (v/v) formic acid (Solvent A) and 100 % (v/v) CH_3_CN in 0.1 % (v/v) formic acid (Solvent B). The flow gradient was (i) 0 - 6 min at 3 % Solvent B, (ii) 6 - 95 min, 3 – 20 % Solvent B (iii) 95 - 105 min, 20 – 40 % Solvent B (iv) 105 - 110 min, 40 – 80 % Solvent B (v) 110 - 115 min, 80 – 80 % Solvent B (vi) 115 - 117 min 85 – 3 % Solvent B and equilibrated at 3% Solvent B for 10 minutes before the next sample injection. In the Data-Dependent Acquisition (DDA) mode, full MS1 spectra were acquired in positive mode, 70 000 resolution from 300 - 1650 *m/z*, AGC target of 3e^6^ and maximum IT time of 50 ms. Fifteen of the most intense peptide ions with charge states ≥ 2 and intensity threshold of 1.7e^4^ were isolated for MSMS. The isolation window was set at 1.2 m/z and precursors fragmented using normalized collision energy of 30, 17 500 resolution, AGC target of 1e^5^ and maximum IT time of 100 ms. Dynamic exclusion was set to be 30 sec. In the Data Independent Acquisition (DIA) mode, the separation gradient was identical to that for DDA analysis. The QExactive plus mass spectrometer was operated in the hyper reaction monitoring/data independent (HRM/DIA) mode, whereby full MS1 spectra were acquired in positive mode from 400 – 1000 *m/z*, 70 000 resolution, AGC target of 3e^6^ and maximum IT time of 50 ms. The isolation window was set at 21 *m/z* with a loop count of 30 and all precursors fragmented using normalized collision energy of 30, 17 500 resolution, AGC target of 1e^6^.

#### Analysis of global proteome of neurons

Three biological replicates per group were used (i.e. n = 3 for control group, n = 3 for each of the treatment groups: treatment with glutamate for 30 min or 240 min). Neuronal lysates (500 µg) were mixed with cold acetone (-20°C) (1:5, v/v) in microfuge tubes and incubated at -20°C overnight to precipitate proteins. Acetone precipitated proteins (in control and treated lysates) were resuspended in 8 M urea in 50 mM TEAB (pH 8.0), and protein estimation was carried out using BCA assay (Pierce-Thermo Scientific) according to manufacturer’s instruction. Equal amounts of protein were reduced with 10 mM tris-(2-carboxyethyl)-phosphine (TCEP) for 45 min at 37°C in a bench top vortex shaker. Reduced samples were alkylated with 55 mM iodoacetamide shaking 45 min at 37°C. Samples were diluted to 1 M urea (diluted with 25 mM TEAB) and digested with sequencing grade modified trypsin (1:50) overnight at 37°C. Digested samples were acidified to 1% (v/v) with pure formic acid, and solid phase extraction (SPE) was carried out with 60 mg Oasis HBL cartridge (Waters) to clean up the digested peptides. Briefly, the cartridge was washed with 80% acetonitrile (ACN) containing 0.1% trifluoroacetic acid (TFA) first and then with only 0.1% TFA before sample loading. Samples were washed again with 0.1% TFA and eluted with 800 µl 80% ACN containing 0.1% TFA. An aliquot (20 µg) of eluted peptides were freeze-dried overnight prior to analysis of changes in global proteome. For quantitative global proteomic analysis, 1 µg peptide in the presence of spiked-in iRT peptide was injected into the mass spectrometer and analysed using the HRM/DIA mode followed by analysis with the Spectronaut DIA-MS methodology and making use of the global proteome-specific spectral library built with the SEQUEST search engine incorporated in Proteome Discover (PD) (Wang, Ma et al., 2020).

#### Analysis of the changes in neuronal N-terminome during excitotoxicity by the Terminal Amine Isotopic labelling of Substrates (TAILS) method

Three biological replicates per group were used (i.e. n = 3 for control group, n = 3 for each of the treatment groups: treatment with glutamate for 30 min or 240 min). Neurons were treated with the neurotoxic concentration of glutamate (100 µM) for 30 min and 240 min. This toxic treatment strategy has been known to induce enhanced limited proteolysis (referred to as proteolytic processing) as well as degradation of specific neuronal proteins by specific proteases activated in response to glutamate over-stimulation (referred to as excitotoxicity-activated proteases) (El-Gebali, Mistry et al., 2019, Hossain et al., 2013). Proteins in the cell lysates of control (untreated) neurons and glutamate-treated neurons were precipitated by ice-cold acetone.

After resuspension and denaturation in 8 M guanidinium hydrochloride in 100 mM HEPES, the neuronal proteins were reduced and alkylated. This is followed by isotopic dimethyl labelling of the free amino groups including the N^α^-amino groups and ε-amino groups of the lysine side chains in the neuronal proteins. Proteins of the control neurons were labelled with formaldehyde (CH_2_O) (referred to as light dimethyl labelled), while those of the glutamate-treated neurons were labelled with deuterated formaldehyde (CD_2_O) (referred to as medium dimethyl labelled). Thus, the neo-N-termini of truncated protein fragments generated by proteolysis catalyzed by the excitotoxicity-activated proteases were medium dimethyl labelled (i.e., covalently linked with deuterated dimethyl groups). The light dimethyl labelled proteins from control neurons and the medium dimethyl labelled proteins from treated neurons were mixed at a ratio of 1:1 (w/w). The protein mixture was subjected to tryptic digestion. Tryptic peptides derived from the N-terminal end of neuronal proteins were devoid of free amino groups because the amino groups were either naturally blocked by modifications such as acetylation and myristoylation *in vivo* or by dimethylation *in vitro*. The other tryptic peptides derived from other parts of neuronal proteins contained the newly formed free N^α^-amino group resulting from trypsinization. These peptides were selectively captured by the Hydroxy Polyglycerol Aldehyde (HPG-ALD) Polymer by reaction of their N^α^- amino groups with the aldehyde (-CHO) groups of the polymer. After ultrafiltration to remove the complex of tryptic peptide-bound HPG-ALD polymers from the naturally modified and dimethyl labelled N-terminal peptides. The modified N-terminal peptides from each sample were pre-fractionated into 4 fractions SDB-RPS (styrene-divinylbenzene reverse phase sulfonate) based fractionation prior to LC-MS/MS analysis on the Orbitrap Elite mass spectrometer. For TAILS analysis of the effects of glutamate treatment in the presence of calpeptin, the same number of animals per group) and the same procedures were used to process neuronal lysates prior to LC-MS/MS analysis.

LC-MS/MS was carried out on LTQ Orbitrap Elite (Thermo Scientific) with a nanoESI interface in conjunction with an Ultimate 3000 RSLC nano HPLC (Dionex Ultimate 3000). The LC system was equipped with an Acclaim Pepmap nano-trap column (Dionex-C18, 100 Å, 75 µm x 2 cm) and an Acclaim Pepmap RSLC analytical column (Dionex-C18, 100 Å, 75 µm x 50 cm). The tryptic peptides were injected to the enrichment column at an isocratic flow of 5 µL/min of 3% v/v CH_3_CN containing 0.1% v/v formic acid for 5 min before the enrichment column was switched in-line with the analytical column. The eluents were 0.1% v/v formic acid (solvent A) and 100% v/v CH_3_CN in 0.1% v/v formic acid (solvent B). The flow gradient was (i) 0-6 min at 3% B, (ii) 6-95 min, 3-20% B (iii) 95-105 min, 20-40% B (iv) 105-110 min, 40-80% B (v) 110-115 min, 80-80% B (vi) 115-117 min 85-3% and equilibrated at 3% B for 10 minutes before the next sample injection. The LTQ Orbitrap Elite spectrometer was operated in the data-dependent mode with nanoESI spray voltage of 1.8kV, capillary temperature of 250°C and S-lens RF value of 55%. All spectra were acquired in positive mode with full scan MS spectra from *m/z* 300-1650 in the FT mode at 240,000 resolution. Automated gain control was set to a target value of 1.0e6. Lock mass of 445.120025 was used. The top 20 most intense precursors were subjected to rapid collision induced dissociation (rCID) with the normalized collision energy of 30 and activation q of 0.25. Dynamic exclusion with of 30 seconds was applied for repeated precursors.

#### Synthesis of FITC-TAT-SRC peptide

Peptides were constructed on a CEM Liberty 12-Channel Automated Microwave Peptide Synthesizer using Fmoc-PAL-PEG-PS^TM^ (Rink resin; loading capacity 0.21 mmol/g) (ThermoFisher, Cat.#: GEN913383) and Fmoc-protected amino acids (GL Biochem (Shanghai)). Fmoc deprotections were performed using 20% piperidine in DMF. Activation of Fmoc-amino acids was achieved using a 0.5 M solution of HCTU (2-(6-chloro-1H-benzotriazole-1-yl)-1,1,3,3-tetramethylaminium hexafluorophosphate) and DIPEA (diisopropylethylamine) in DMF in a ratio of 2 ml:2 ml:0.34 ml per 1 mmol of Fmoc amino acid used. Peptide coupling and deprotection efficiency was monitored using a 2,4,6-trinitrobenzene-sulfonic acid (TNBSA) assay (Hancock & Battersby, 1976). The FITC-TAT-ahx-SRC(49-79) peptide (FITC-Tat-Src) was synthesized sequentially as two main parts with aminohexanoic acid linker used between the two main sequences. First, the sequence ahx-SRC 49-79 was synthesized. The synthesized peptide was validated via ESI-MS and RP-HPLC. Then, the TAT peptide with sequence ahx-GRKKRRQRRRPQ was continued on the ahx-SRC 49-79 sequence while still on resin to generate ahx-TAT-ahx-SRC(49-79) peptide-resin. Fluorescein isothiocyanate (FITC) was coupled to ahx-TAT-ahx-SRC(49-79) peptide-resin using HCTU and DIPEA. The peptide was cleaved from the resin using TFA/triisopropylsilane/H_2_O mixture (volume ratio of 95:2.5:2.5) for 90 min. Excess TFA was removed via evaporation with stream of N_2_ gas. The peptide was precipitated by addition of diethyl ether. The mixture was then centrifuged and the ether decanted. The pellet containing the peptide was re-dissolved in 30% acetonitrile/H_2_O and filtered through a 0.22 μm filter. The crude peptide solution was lyophilized prior to purification by semi-preparative RP-HPLC. Fractions containing the pure peptide were pooled and lyophilized. The dried peptide was stored at 4 °C until further use. The purified peptide was analyzed and validated via ESI-MS with mass [M + H^+^] of 5385.4 Da and RP-HPLC shown as a single peak in HPLC chromatogram.

### Quantification and statistical analysis of proteomics data

#### Quantitation of global proteomic changes in mouse primary cortical neurons induced by glutamate treatment

Protein/peptide identification from the DDA-based analysis and subsequent spectral library generation were conducted using Proteome Discoverer (v.2.1, Thermo Fischer Scientific) with the Sequest HT search engine in combination with the Percolator semi-supervised learning algorithm (Kall, Canterbury et al., 2007) and PhosphoRS (Taus, Kocher et al., 2011) or with MaxQuant software utilizing the Andromeda search engine (Tyanova, Temu et al., 2016) on M*us musculus* protein database (SwissProt (TaxID = 10090)(version 2017-10-25)) (Figure S14). The search parameters were: a precursor tolerance of 20 ppm, MSMS tolerance of 0.05 Da, fixed modifications of carbamidomethylation of <u>cysteine</u> (+57 Da), <u>methionine</u> oxidation (+16 Da) or phosphorylation of serine, threonine and tyrosine (+80Da) . Peptides were accepted based on a FDR of <0.01 at both the <u>peptide and protein</u> levels.

DIA based quantitative analysis was carried out using the Spectronaut software (Spectronaut 11, Biognosys). To build the spectral library in Spectronaut based on MaxQuant and Proteome Discoverer results, all the default parameters were used except that best N-terminal fragments per peptide was set to minimum 5 and maximum 20. Two spectral libraries were built for phosphopeptides whereas one library from the search results of Proteome Discoverer was built for the global library. The spectral libraries for peptides are generated identically (Figure S14). The DIA files for each sample were converted to htrms format sing HTRMS Converter (Biognosy) and loaded on Spectronaut 11 for generation of protein intensities. The default parameters were used for data extraction, XIC extraction, calibration, identification, and protein inference. The iRT profiling strategy was used with unify peptide peak options enabled. For quantitation of the trptic peptides unique to each protein, the Q value percentile of 0.75 was set. Quantitation is based on stripped sequence, and global normalization was performed. The Spectronaut results output were further processed with Perseus software (version 1.6) (Tyanova & Cox, 2018).

#### Key resources

The antibodies used in our studies include: rabbit polyclonal anti-CRMP2 antibody (RRID: AB_2094339; Cat. No.: 9393; Cell Signaling Technology), rabbit polyclonal anti-CRMP2 (phospho T509) antibody (Cat. No.: ab192799; Abcam), anti-DCLK1 (anti-DCMKL1) antibody against amino acids 679-729 near the C-terminus of DCLK1 (Cat. No.: ab106635; Abcam), mouse anti-Src Mab327 antibody (RRID: AB_443522; Cat. No.: ab16885; Abcam), anti-CD11b (Clone OX-42) mouse monoclonal antibody (Cat. No.: MCA275G; BioRad (formerly serotec)), NeuN rabbit polyclonal antibody (RRID: AB_10807945; Cat. No.: ABN78; MERK Millipore (Sigma Aldrich)), anti-GFAP mouse monoclonal antibody (Clone GA5) (RRID: AB_11212597; Cat. No.: MAB360; MERK Millipore (Sigma Aldrich), CaMKIIα (#NB100-1983, RRID:AB_10001339; mouse monoclonal IgG, clone 6g9, Novus Biologicals), CaMKII (pan) (#4436, RRID:AB_1054545; rabbit monoclonal IgG, clone D11A10, Cell Signaling Technology), beta-Actin (#ab6276, RRID: AB_2223210; mouse monoclonal IgG, cloneAC-15, Abcam), CaMKIIβ (#ab34703, RRID:AB_2275072; rabbit polyclonal antibody, Abcam); Goat anti-mouse Alexa Fluor Plus 800 (#A32730, RRID: AB_2633279; polyclonal IgG, Invitrogen) and donkey anti-rabbit Alexa Fluor Plus 488 (#A32790, RRID: AB_2762833; polyclonal IgG, Invitrogen).

The following recombinant proteins were used: CaMKIIα (#PR4586C, Invitrogen), GST-CaMKIIβ (#02-110, Carna biosciences) and Calpain-1 (#208713-500UG, Calbiochem).

Chemicals and reagents specifically used for proteomic analysis include: Pierce™ high pH reversed-phase peptide fractionation kit (Cat. No.: 84868; ThermoFisher), High-Select™ Fe-NTA phosphopeptide enrichment kit (Cat. No.: A32992; ThermoFisher), titansphere phos-TiO_2_ (Cat. No.: 5010-21315; GL Sciences), triehylammonium bicarbonate (TEAB) buffer (Cat. No.:5010-21315; Sigma), formaldehyde (DLM-805-PK (CD_2_O) ULM-9498-PK (CH_2_O), Cambridge Isotope) and hydroxy polyglycerol aldehyde (HPG-ALD) polymer (UBC.FLINTBOX https://ubc.flintbox.com/#technologies/888fc51c-36c0-40dc-a5c9-0f176ba68293).

The chemicals used for synthesis of the cell permeable TAT-Src, TAT-Scrambled and fluorescent TAT-Src peptides include: Fmoc-PAL-PEG-PS^TM^ (Rink resin; loading capacity 0.21 mmol/g) (Cat. No.: GEN913383; ThermoFisher), Fmoc-protected amino acids (GL Biochem (Shanghai)), phalloidin–tetramethylrhodamine B isothiocyanate (Phalloidin-Tritc) (Cat. No.: Sigma P1951; Merck).

The animals used in our studies including (i) C57BL/6 mice (pregnant and at gestational day 14-15) used for cultured primary cortical neurons (ii) C57BL/6 mice used for construction of spectral libraries and (iii) Male hooded Wistar rats for the in vivo model of neurotoxicity were sourced from Animal Resources Centre, Canning Vale WA 6970 Australia (iv) male C57BL/6*J* were obtained from the Biomedical Research Facility, University of Otago, New Zealand and *Camk2a^-/-^* mice (*Camk2*^atm3Sva^, MGI:2389262) mice backcrossed into the C57BL/6*J* background were bread in the In Vivo Pharmacology Research Unit, University of Copenhagen, Denmark for the use in photothrombotic stroke surgeries.

The software and algorithms used in our studies are: Proteome Discoverer (Thermo Scientific; RRID: SCR_014477), MaxQuant (RRID: SCR_014485), Spectronaut (Spectronaut 11) (Biognosys), PhosphoSitePlus: Protein Modification Site (Cell Signaling Technology; RRID: SCR_001837), Image J (RRID: SCR_003070), QIAGEN Ingenuity Pathway Analysis (Qiagen) and SynGO (RRID:SCR_017330), the protein interaction resource derived from cross-linking mass spectrometry database of synaptosome and microsome fractions purified from hippocampus and cerebellum of mouse brains (Gonzalez-Lozano et al., 2020) and String database (Szklarczyk et al., 2019)

## Supporting information

Supplemental Results and Figures S1-S12

## Acknowledgments

We thank Robert Qi, Swati Varshney and Anderly Chueh for comments and suggestions for drafting this manuscript.

## Author Contributions

S.S.A. and C.-S.A. conceptualized, designed, performed, and analyzed most of the experiments and co-wrote the paper. H.-C.C., N.A.W., A.D., O.K., G.D.C. and C.R. conceptualized, designed and analyzed the proteomic data, neuronal cell biology data or data of in vivo model of neurotoxicity. M.I.H., M.A.K., H.-J.Z., G.D.C., A.H., L.B. and C.R. conceptualized, designed and performed the experiments of excitotoxic cell death of cultured neurons and effects of Tat-Src and Tat-Scrambled peptides on neuronal loss in vivo. N.G.-K. A.N.C and P.W. contributed to generation and interpretation of *in vitro* and *in vivo* data of CaMKII. S.S.A., H.-C.C., C.-S.A., N.A.W., S.S., H.N., R.M. and D.D. performed experiments to generate the spectral libraries for DIA proteomic analysis of the changes in global and phospho-proteomes of neurons. M.L., H.-C.C., A.D., O.K. N.A.W. and C.-S.A. analyzed the N-terminomic data to define the potential substrates of calpains in neurons. H.-C.C. and P.P. performed the SynGO analysis to assign the synaptic locations and biological processes of the significantly modified proteins. D.L., H.-J.Z., A. Dhillon, A. Dufour, I.S.L. P.P. F.B. A.N.C. and J.P.L. conceptualized, interpreted experimental data on excitotoxic neuronal death and experimental validation of the proteomic results and co-wrote the paper.

## Data and materials availability

The mass spectrometry proteomics data have been deposited to the ProteomeXchange Consortium via the PRIDE (Perez-Riverol, Csordas et al., 2019) partner repository with the dataset identifiers PXD019211 for the project entitled “N-terminomics Analysis of Mouse cultured cortical neuron treated with glutamate”, and PXD019527 for the project entitled “Phosphoproteomics of mouse cortical neuron”.

