## Supplemental Results and Figures S1-S12 for "N-Terminomic Changes of Neurons During Excitotoxicity Reveal Proteolytic Events Associated with Synaptic Dysfunctions and Inform Potential Targets for Neuroprotection"

**This file includes** *Supplementary Results and Figures. S1 to S12*

**Other Supplementary Materials for this manuscript include** *Tables S1 to S9 (in excel files)*

| | |
|---|---|
| Table S1 | TAILS_ Glutamate 30min/Control (Separate file) |
| Table S2 | TAILS_ Glutamate 240min/Control (Separate file) |
| Table S3 | Global/Shotgun Proteomics DIA (Separate file) |
| Table S4 | Abundance ratios of neo-N-terminal peptides and unique peptides of significantly proteolyzed proteins in excitotoxic neurons (Separate file) |
| Table S5 | TAILS_ Glutamate + Calpeptin 30min/Control (Separate file) |
| Table S6 | TAILS_ Glutamate + Calpeptin 240min/Control (Separate file) |
| Table S7 | Comparison of Significantly cleaved Peptides between Glutamate and Glutamate + Calpeptin (Separate file) |
| Table S8 | Annotated Table for Functional Domain_TAILS (Separate file) |
| Table S9 | SynGO analysis of neuronal proteins undergoing significant proteolytic processing during excitotoxicity (Separate file) |

### Supplementary Results

*Classification of the N-terminal peptides identified and quantified by TAILS method*

Using the TAILS method (Kleifeld et al., 2011), we identified and quantified (i) the "natural" free N-termini of mature proteins biosynthesized in physiological conditions, and (ii) the newly formed neo-N-termini of stable truncated protein fragments and intermediate peptide fragments generated by proteolysis of neuronal proteins during excitotoxicity. Of the quantifiable N-terminal peptides identified by the TAILS analysis of excitotoxic neurons, over 2,300 contained acetylated N-termini and over 2,600 contained dimethyl labelled N-termini (Figure S2A, Tables S6A and S7A). Whereas the acetylated N-terminal peptides were derived from intact proteins undergoing N-terminal acetylation during biosynthesis and maturation in neurons, the dimethyl-labelled N-terminal peptides were derived from free N-termini of neuronal proteins *in vitro* via the TAILS procedure. The dimethyl labelled N-terminal peptides were further sub-divided into four groups: (i) those retaining the first methionine encoded by the start codon (+ Met); (ii) those with the start codon-encoded methionine removed (-Met); (iii) those with the N-terminal signal peptide segments removed during maturation of the parental neuronal proteins *in vivo* (-signal peptide); and (iv) those containing neo-N-termini generated by proteolysis of intact mature neuronal proteins (Neo) (Figure S2A). We focused our further analysis on the neo-N-terminal peptides as they revealed both the cleavage sites and identities of neuronal proteins undergoing proteolysis during excitotoxicity. Based on the rationale depicted in Figure S2C, neo-N-terminal peptides exhibiting increased abundance were assigned as those derived from the stable truncated protein fragments generated from enhanced proteolytic processing of neuronal proteins during excitotoxicity. Neo-N-terminal peptides exhibiting decreased abundance were assigned as being derived from neuronal proteins undergoing enhanced degradation for clearance during excitotoxicity (Figure S2B).

*Determination of the abundance ratio cut-off values to identify neuronal proteins exhibiting significantly enhanced proteolysis induced by treatment with glutamate or co-treatment with glutamate and calpeptin*

Over 2,000 neo-N-terminal peptides were found by the TAILS method to be generated by enhanced proteolysis during excitotoxicity (Figure S2A). Figures S3 and S7 depict results of our statistical analysis to calculate the thresholds for their assignment to be neo-N-terminal peptides generated by significantly enhanced degradation and those generated by significantly enhanced proteolytic processing of neuronal proteins during excitotoxicity. Based upon the normalized distribution of the $\log_2$ M/L ratios of the identified neo-N-terminal peptides, we determined the median and standard deviations (S.D.) of the distribution of the $\log_2$ M/L ratios. Statistically, values that are outside the 1.5 interquartile range (IQR) are considered as outliers (Tukey, 1977) (Figure S3A). The 1.5×IQR values can be calculated as 5 × S.D. of the distributions of the abundance ratios ($\log_2$ M/L ratios) of all quantifiable N-terminal peptides (Figure S3B). Using the Tukey's 1.5 × interquartile range (IQR) rule (Tukey, 1977), we determined the abundance ratio cut-off values (M/L) for the neo-N-terminal peptides exhibiting significantly reduced abundance and those exhibiting significantly increased abundance during excitotoxicity (Figure S3A).

To determine the abundance ratio cut-off values to identify the neo-N-terminal peptides generated by significant proteolysis by neuronal proteases activated in response to co-treatment with glutamate and calpeptin, the M/L ratio cut-off values equivalent to 1.5 interquartile range (IQR) were calculated. As depicted in Figure S7A, for the N-terminal peptides co-treated for 30 min, the lower and upper cut-off values were 0.271 and 3.8 (i.e. $M/L \geq 3.8$ for the neo-N-terminal peptides generated by significantly enhanced proteolytic processing and $M/L \leq 0.271$ for those generated by significantly enhanced degradation). For those derived from neurons co-treated for 240 min, the lower and upper cut-off values were 0.386 and 2.876, respectively.

### Supplementary Figures S1-S12

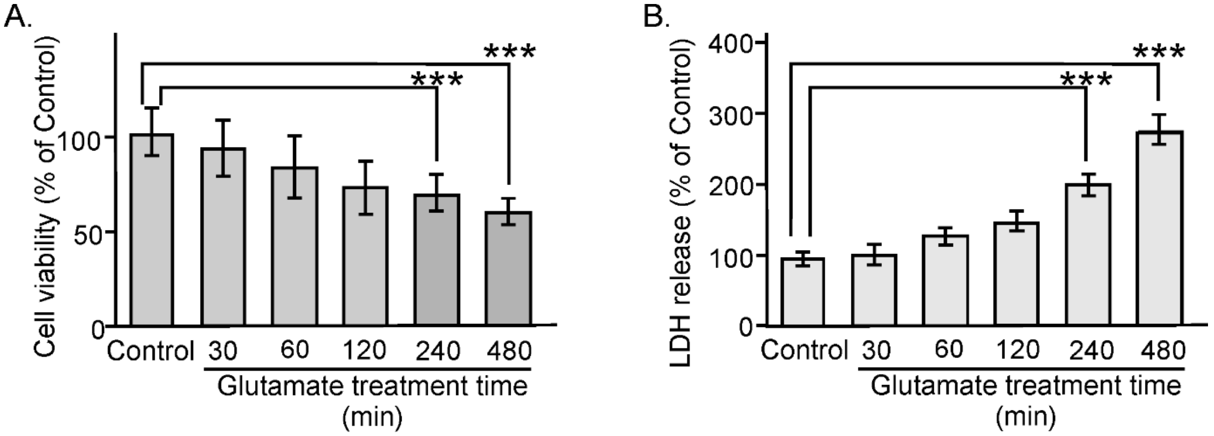


**Fig. S1 Viability of cultured cortical neurons after different duration of glutamate treatment**

**A.** MTT assay results showing neuronal viability at different time points after glutamate treatment. The amount of formazon formed by the viable neurons at each time point of treatment is presented as the percentage of that formed by the viable untreated neurons (Control). Data represent as mean ± S.D. (error bars), n = 6; *** represents $p < 0.005$, Student's *t*-test. **B.** The amount of LDH released from the damaged neurons at each designated treatment time point is presented as the percentage of that released by the untreated neurons (Control). Data represent as mean ± S.D. (error bar), n = 3; *** represents $p < 0.005$, Student's *t*-test.

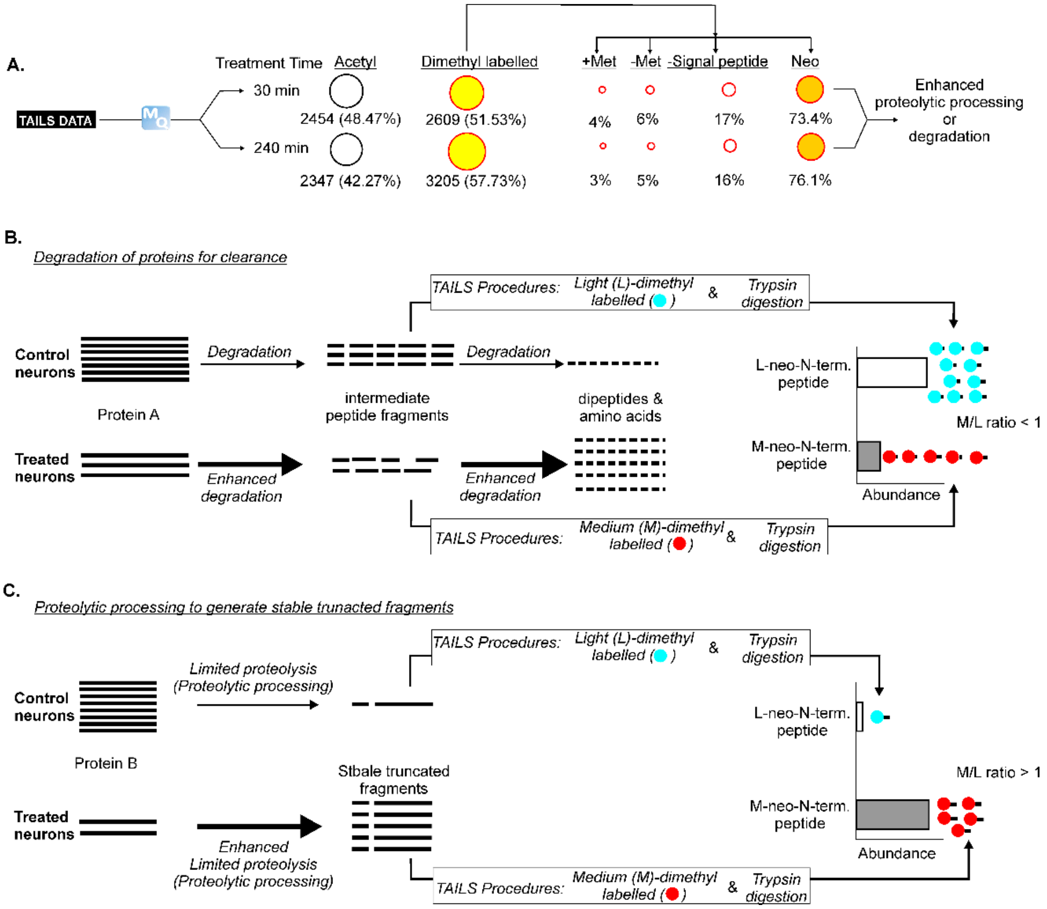


**Figure S2 Classification of identified N-terminal peptides derived from neuronal proteins**

**A.** The search engine Andromeda of MaxQuant (MQ) was used to identify and quantify the N-terminal peptides derived from tryptic digests of the neuronal cell lysates in the TAILS method. The identified N-terminal peptides were classified into those with acetylated N-termini and those with free N-termini which were isotopically dimethyl labelled. The dimethyl-labelled N-terminal peptides were sub-classified into those (i) with intact N-terminal methionine encoded by the start codon (+Met), (ii) with the N-terminal methionine post-translationally deleted *in vivo* (-Met), (iii) with the signal peptide sequence post-translationally deleted by signal peptidases *in vivo* (-Signal peptide) and (iv) with neo-N-termini generated by enhanced proteolysis of neuronal proteins during excitotoxicity (Neo). **B.** Assignment of neo-N-terminal peptides generated by enhanced degradation. The treatment induces a hypothetical protein (Protein A) to undergo enhanced degradation for clearance. It is first proteolyzed to form intermediate peptide fragments, which are then further degraded to form dipeptides and amino acids. In the TAILS procedures, the intermediate peptide fragments are isotopically dimethyl labelled at their neo-N-termini, followed by tryptic digestion to generate the isotopically labelled neo-N-terminal peptides. Since the treatment induces enhanced degradation of the intermediate peptide fragments, the neo-N-terminal peptides derived from Protein A are less abundant in the treated neurons than those in the control neurons (i.e. M/L ratio <1). **C.** Assignment of neo-N-terminal peptides derived from stable protein fragments generated by enhanced proteolytic processing of neuronal proteins. The treatment induces a hypothetical protein (Protein B) to undergo enhanced limited proteolysis (proteolytic processing) to form stable truncated protein fragments. Upon isotopic dimethyl labelling of the neo-N-termini of the truncated protein fragments and tryptic digestion in the TAILS procedures, the resultant labelled neo-N-terminal peptides derived from Protein B are more abundant in the treated neurons than those in the control neurons (i.e. M/L ratio <1).

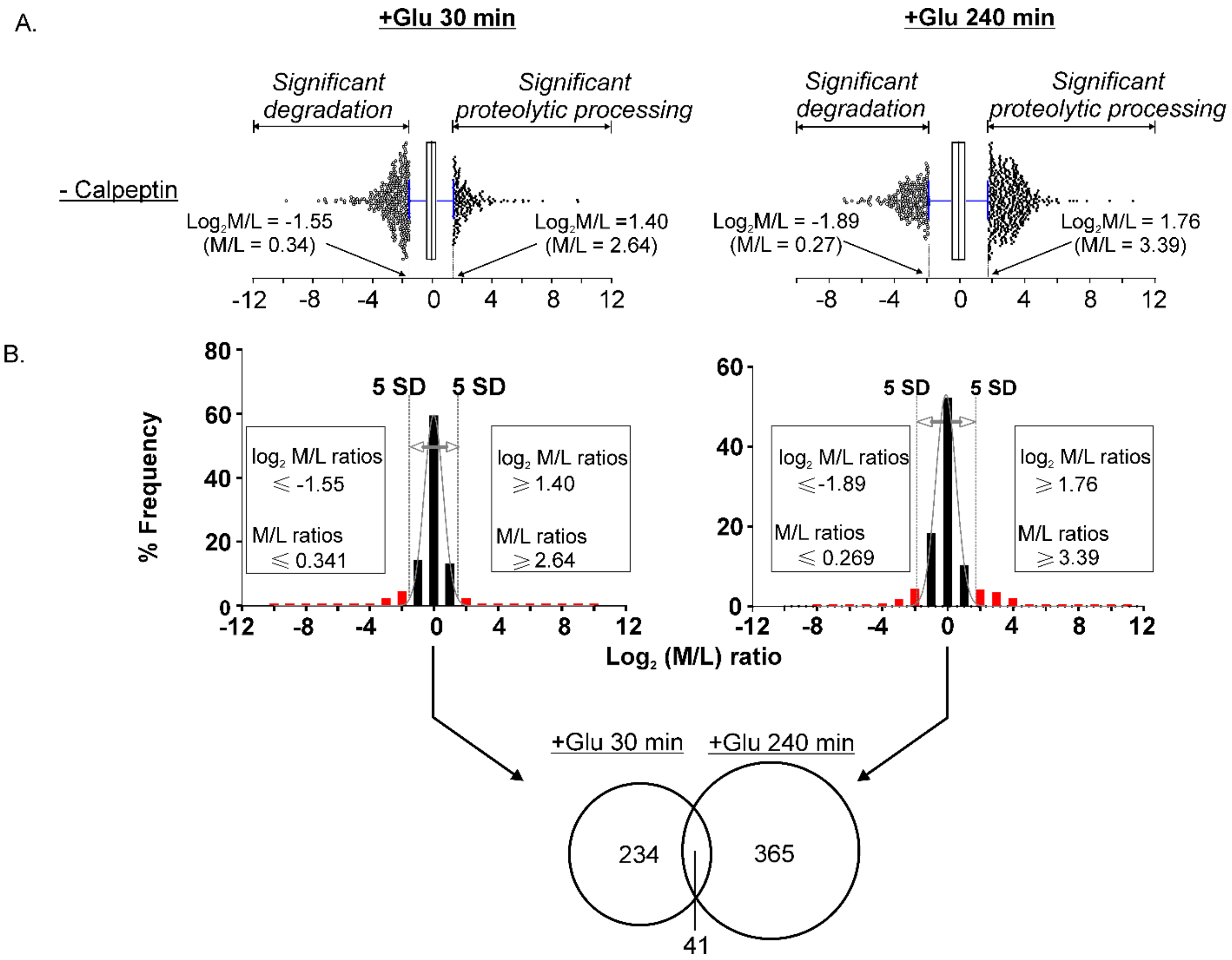


**Figure S3 Determination of the abundance ratio (M/L) cut-off values for classification of neo-N-terminal peptides with significantly altered abundance during excitotoxicity.**

**A.** Box-and-whisker plots of M/L ratio cut-off values determining if the neo-N-terminal peptides were generated by significantly enhanced degradation or by significantly enhanced proteolytic processing. **B.** Histograms showing normal distribution of the data after 30 and 240 min of glutamate treatment. The M/L ratio cut-off values to define if the neo-N-terminal peptides were derived from proteins undergoing significantly enhanced degradation or proteolytic processing during excitotoxicity are indicated. Lower panel: With these cut-off values, 234 and 365 neo-N-terminal peptides were derived from proteins undergoing significant proteolysis in neurons treated with glutamate at 30 min and 240 min, respectively. Among them, 41 were from neurons at both treatment time points (Venn diagram).

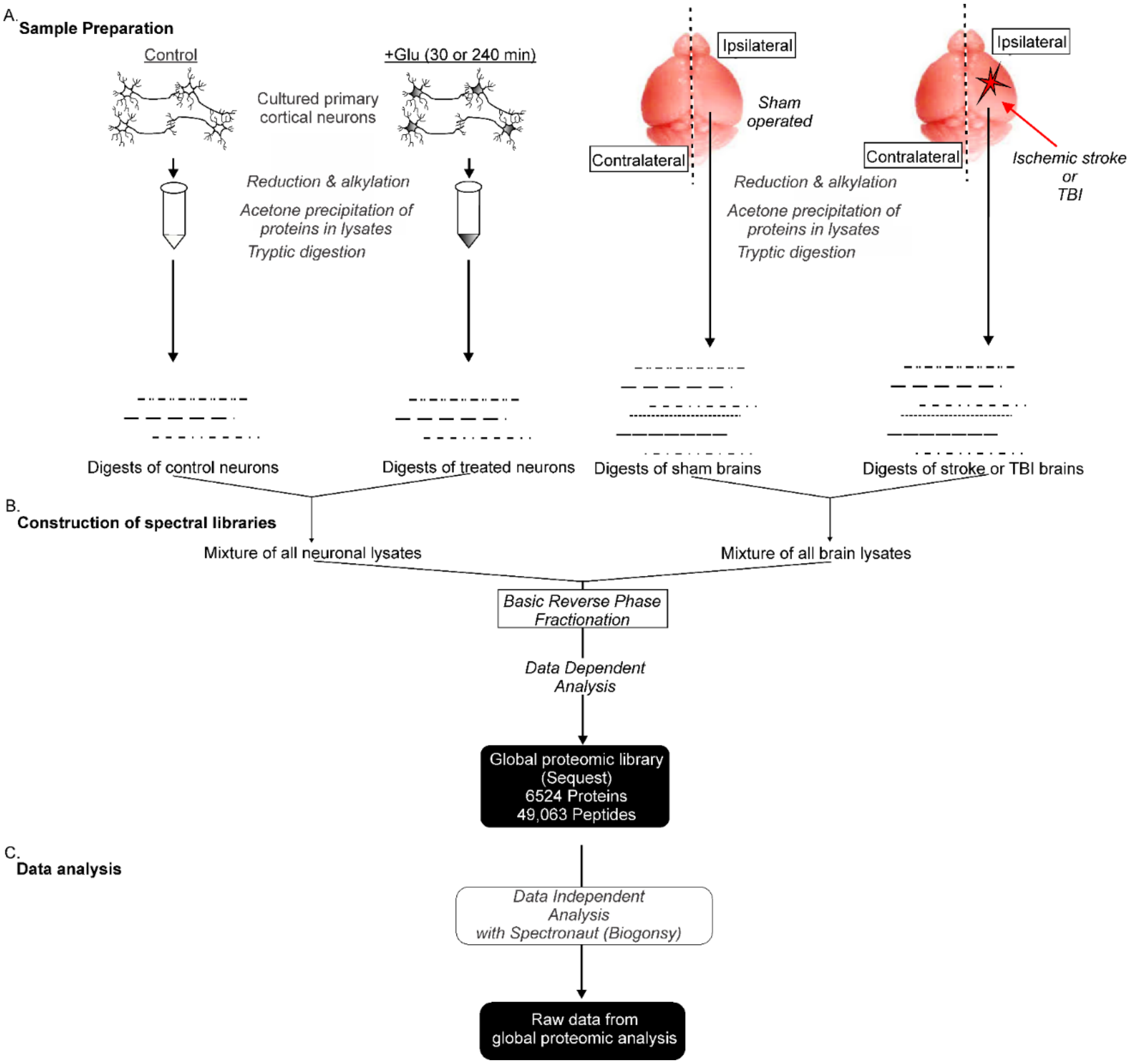


**Figure S4 Workflow for the construction of spectral libraries from lysates of mouse neurons and brain tissues for DIA global proteomic analysis**

**A.** Lysates of untreated (Control) and glutamate-treated neurons were pooled and combined with lysates of brain tissues from sham operated mice and from mouse models of ischemic stroke and traumatic injury (TBI). Proteins in the combined cell and tissue lysates were precipitated, resuspended, reduced and alkylated prior to tryptic digestion. **B.** The resulting tryptic peptides were purified by solid phase extraction (SPE) (Oasis HBL cartridge, Waters) followed by fractionation with basic reverse-phase column chromatography. An aliquot consisting of 20 µg of peptides in each fraction were subjected to LC-MS/MS with the QE plus Orbitrap mass spectrometer for the construction of the spectral library for global proteomic analysis. The identified peptides were used for construction of the spectral libraries. **C.** Data independent analysis (DIA) to profile changes in global proteome of cortical neurons induced by glutamate treatment.

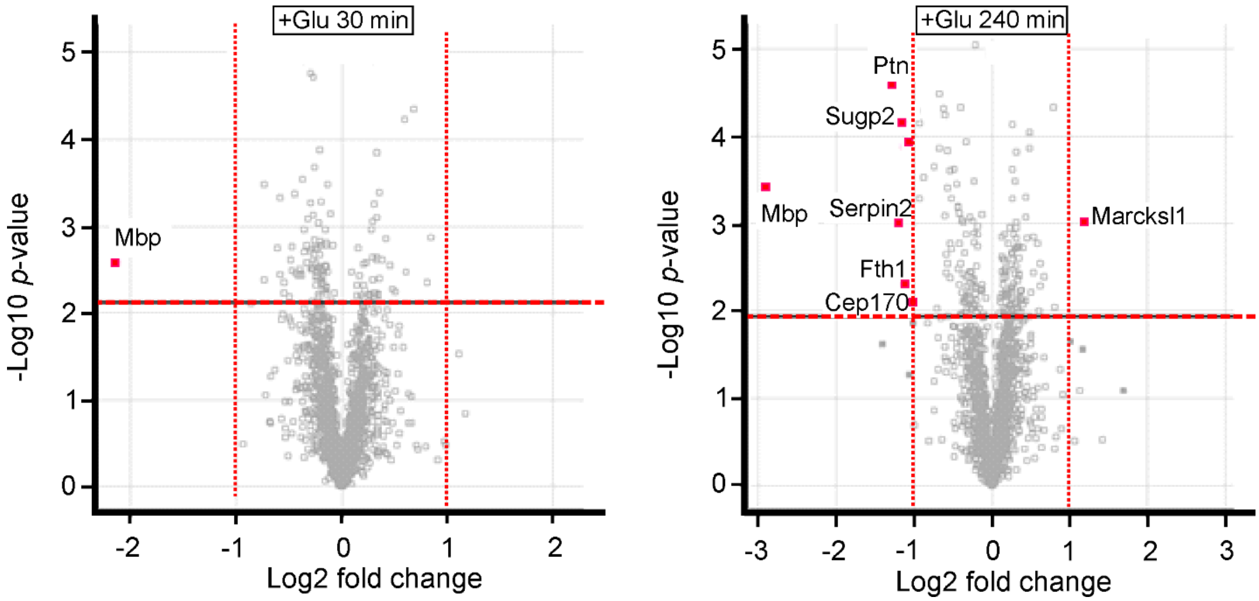


**Figure S5 Global proteomic analysis revealed only a few neuronal proteins exhibiting significant changes in abundance at 30 and 240 min after glutamate treatment**

Volcano plots showing abundance ratios of the identified proteins in glutamate-treated versus Control neurons. The red dotted line in each plot indicates the threshold of false discovery rate (FDR) ≤ 5% (or 0.05) in the two-sample t-test. The fold change for each protein was calculated as $\log_2$ (Treatment/Control abundance) ratio. Red dots depict proteins exhibiting significantly changed abundance (presented as Uniprot accession numbers) with FDR ≤ 0.05 and Treated/Control abundance ratio ≥ 2 or ≤ 0.5. The identified proteins that did not exhibit significant abundance change or with FDR ≥ 0.05 are represented as grey colored square boxes.

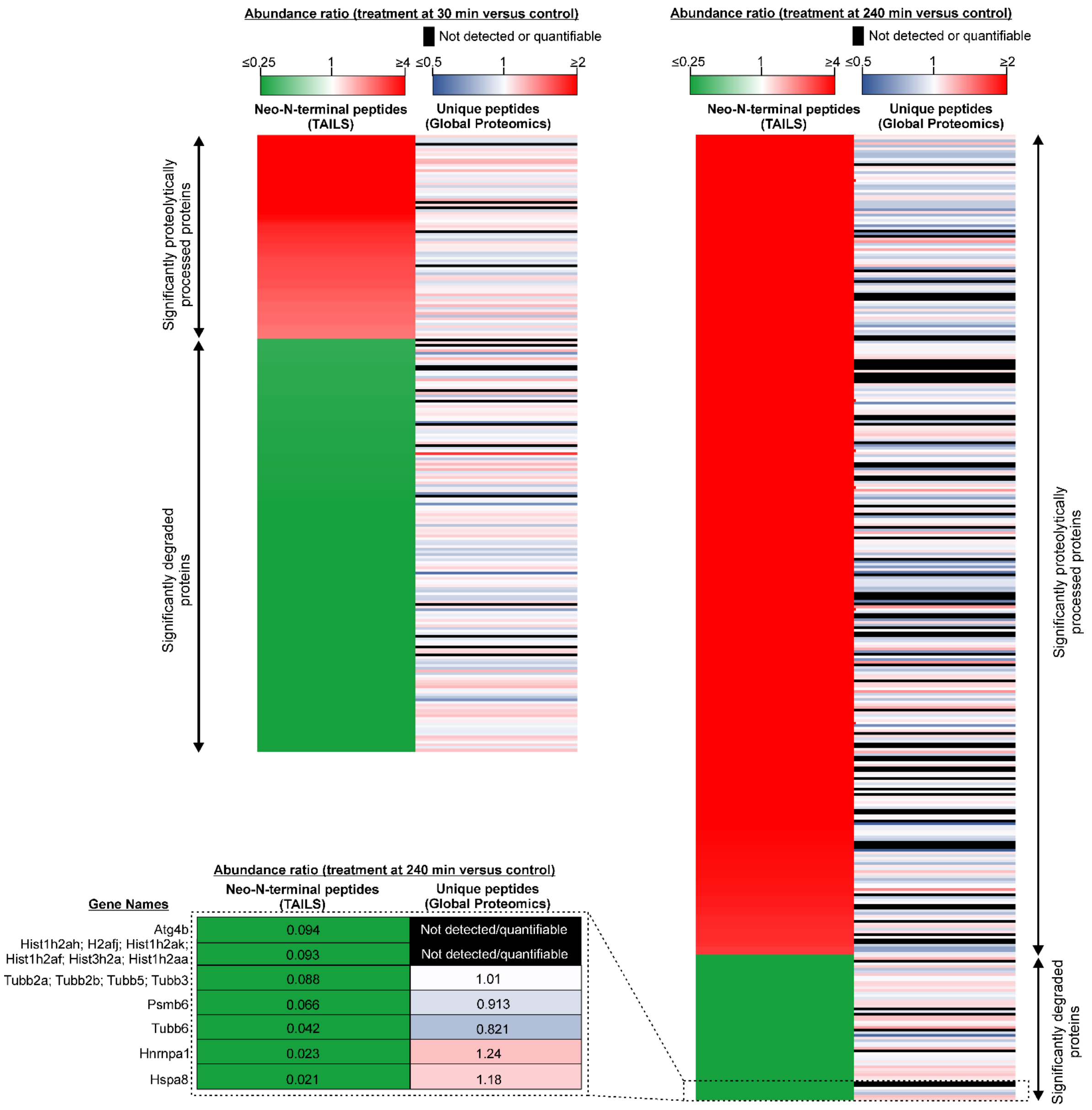


**Figure S6 Abundance ratios of neo-N-terminal peptides and those of the corresponding neuronal proteins undergoing significant proteolysis during excitotoxicity**

Heatmaps depicting the abundance ratios of neo-N-terminal peptides and unique peptides of neuronal proteins undergoing significant proteolytic processing and degradation during excitotoxicity. Inset: A section of the heatmaps and the associated abundance ratio values of the selected neuronal proteins undergoing significant degradation after 240 min of glutamate treatment as depicted in Table S4B.

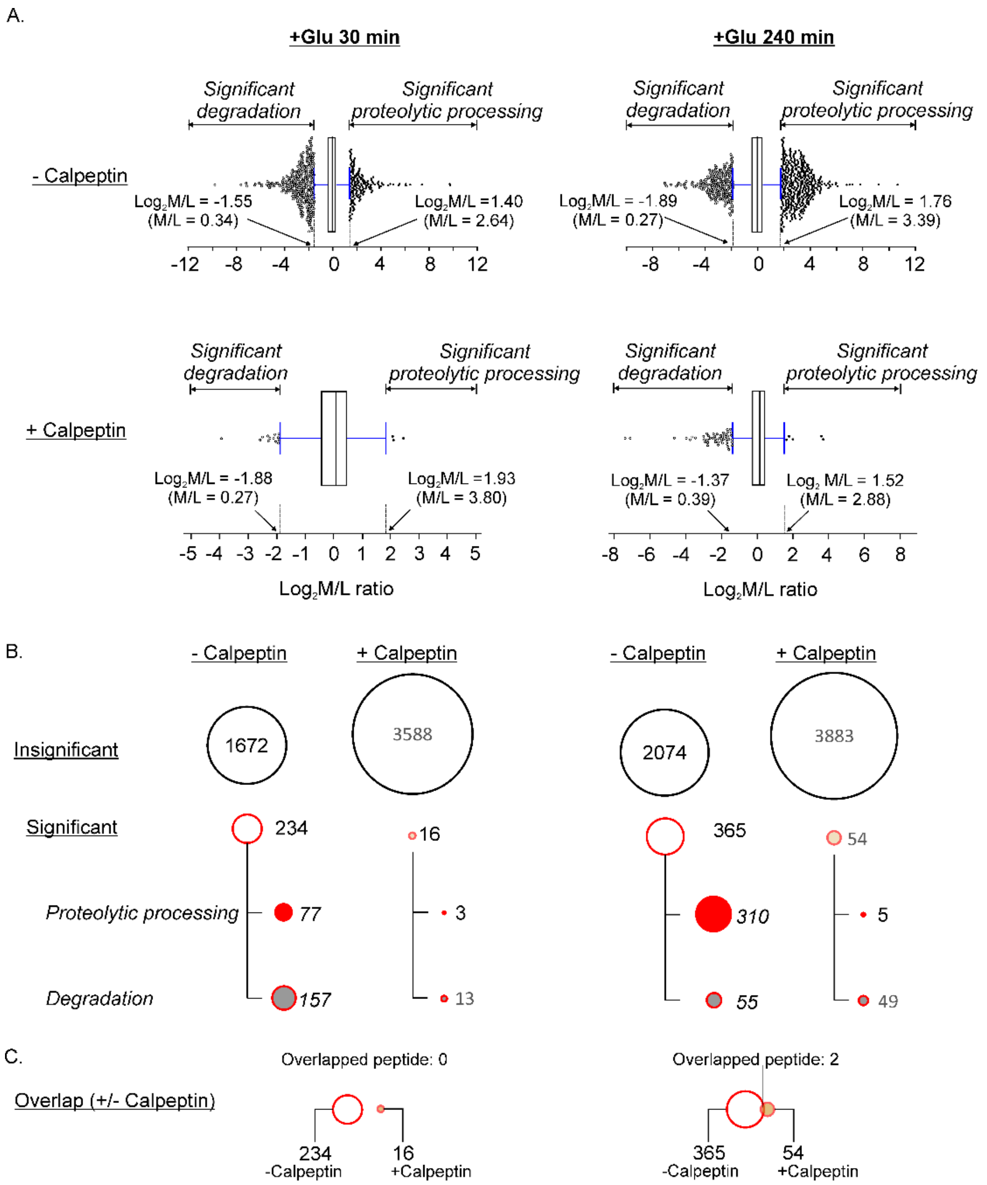


**Figure S7 Neo-N-terminal peptides derived from neuronal proteins induced by glutamate treatment with and without calpeptin**

**A.** Box-and-whisker plots (upper panels) of M/L ratio cut-off values determining if the neo-N-terminal peptides were generated by significantly enhanced degradation or by significantly enhanced proteolytic processing in neurons treated with glutamate for 30 min and 240 min (+Glu 30 min and +Glu 240 min) in the presence and absence of calpeptin (+/- Calpeptin). The M/L ratio cut-off values to define if the neo-N-terminal peptides were derived from proteins undergoing significantly enhanced degradation or proteolytic processing during excitotoxicity. The statistical method to determine these cut-off values is illustrated in Figure S3. **B.** Bubble plots depicting the number of neo-N-terminal peptides exhibiting significant changes in abundance (Significant) and those without significant changes in abundance (Insignificant) in excitotoxicity determined by the M/L cut-off values. The "significant" group of peptides were subdivided into neo-N-terminal peptides derived from proteins undergoing significantly enhanced proteolytic processing and those derived from proteins undergoing significantly enhanced degradation. C. Overlaps of neo-N-terminal peptides derived from proteins undergoing significant proteolysis in neurons treated with glutamate for 30 min and 240 min in the presence and absence of calpeptin.

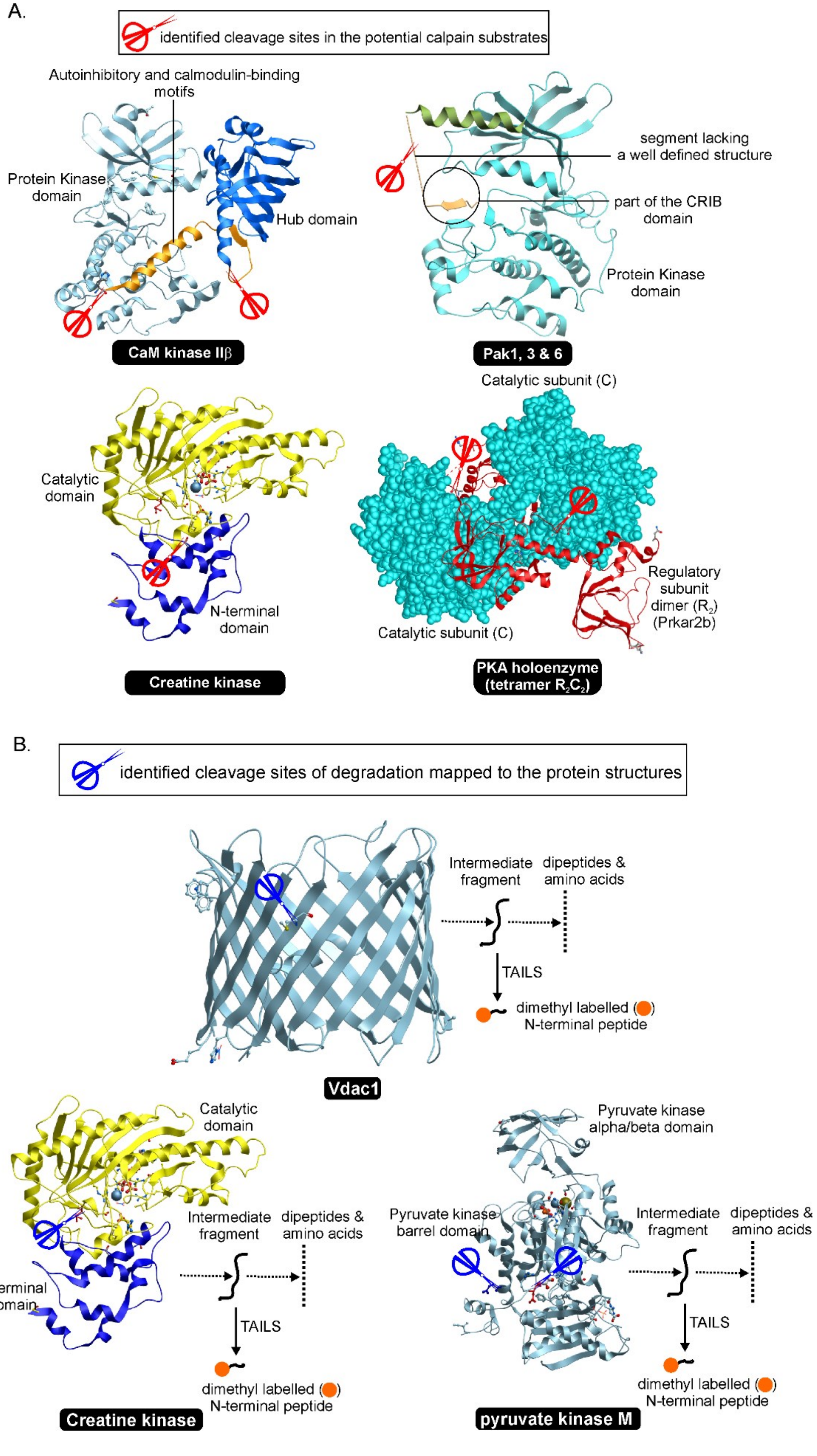
A.
identified cleavage sites in the potential calpain substrates
Autoinhibitory and calmodulin-binding motifs
Protein Kinase domain
Hub domain
CaM kinase IIβ
segment lacking a well defined structure
part of the CRIB domain
Protein Kinase domain
Pak1, 3 & 6
Catalytic domain
N-terminal domain
Creatine kinase
Catalytic subunit (C)
Regulatory subunit dimer ($R_2$) (Prkar2b)
Catalytic subunit (C)
PKA holoenzyme (tetramer $R_2C_2$)
B.
identified cleavage sites of degradation mapped to the protein structures
Intermediate fragment
dipeptides & amino acids
TAILS
dimethyl labelled ( ) N-terminal peptide
Vdac1
Catalytic domain
N-terminal domain
Intermediate fragment
dipeptides & amino acids
TAILS
dimethyl labelled ( ) N-terminal peptide
Creatine kinase
Pyruvate kinase alpha/beta domain
Pyruvate kinase barrel domain
Intermediate fragment
dipeptides & amino acids
TAILS
dimethyl labelled ( ) N-terminal peptide
pyruvate kinase M

**Figure S8 Cleavage sites identified in selected proteins in neurons during excitotoxicity**

**A.** Proteins undergoing enhanced proteolytic processing catalyzed by calpains. Calmodulin-dependent protein kinase IIβ (CaM kinase IIβ), Creatine kinase, p21-activated kinases 1, 3 and 6 (PAK1, 3, & 6) and the regulatory subunit ($R_2$) of cAMP-dependent protein kinase (Prkar2b) were identified by the TAILS method to underwent significantly enhanced proteolytic processing during excitotoxicity. The ribbon model representations of their three dimensional structures are depicted. The red scissors indicate the locations of the identified cleavage sites. The PDB accession codes of these structures are: CaM kinase IIβ, 3SOA; PAK1,3 & 6, 3tnp (this is the structure of PAK4, which shows a high degree of sequence homology with PAK1, 3 and 6); creatine kinase, 3b6r; holoenzyme of cAMP-dependent protein kinase (PKA) with Prkar2b as the regulatory subunits ($R_2$) and two catalytic subunits (C subunit) with the protein kinase domain, 3tnp. All cleavage sites in these proteins are not located within a functional domain; they are located in linker regions, which adopt either a flexible loop structure or a disordered structure. **B.** Proteins undergoing enhanced degradation. Voltage-dependent anion-selective channel protein 1 (Vdac1), Creatine kinase and pyruvate kinase M were identified by the TAILS method as neuronal proteins undergoing significantly enhanced degradation for clearance in excitotoxicity. The red scissors indicate the locations of the identified cleavage sites. As these sites are within the properly folded proteins or functional domains, the identified neo-N-terminal peptides were likely generated by enhanced cleavage of the intermediate fragments in neurons during excitotoxicity. The PDB accession codes of these structures are: Vdac1, 6g6u; creatine kinase, 3b6r; pyruvate kinase M, 3srf.

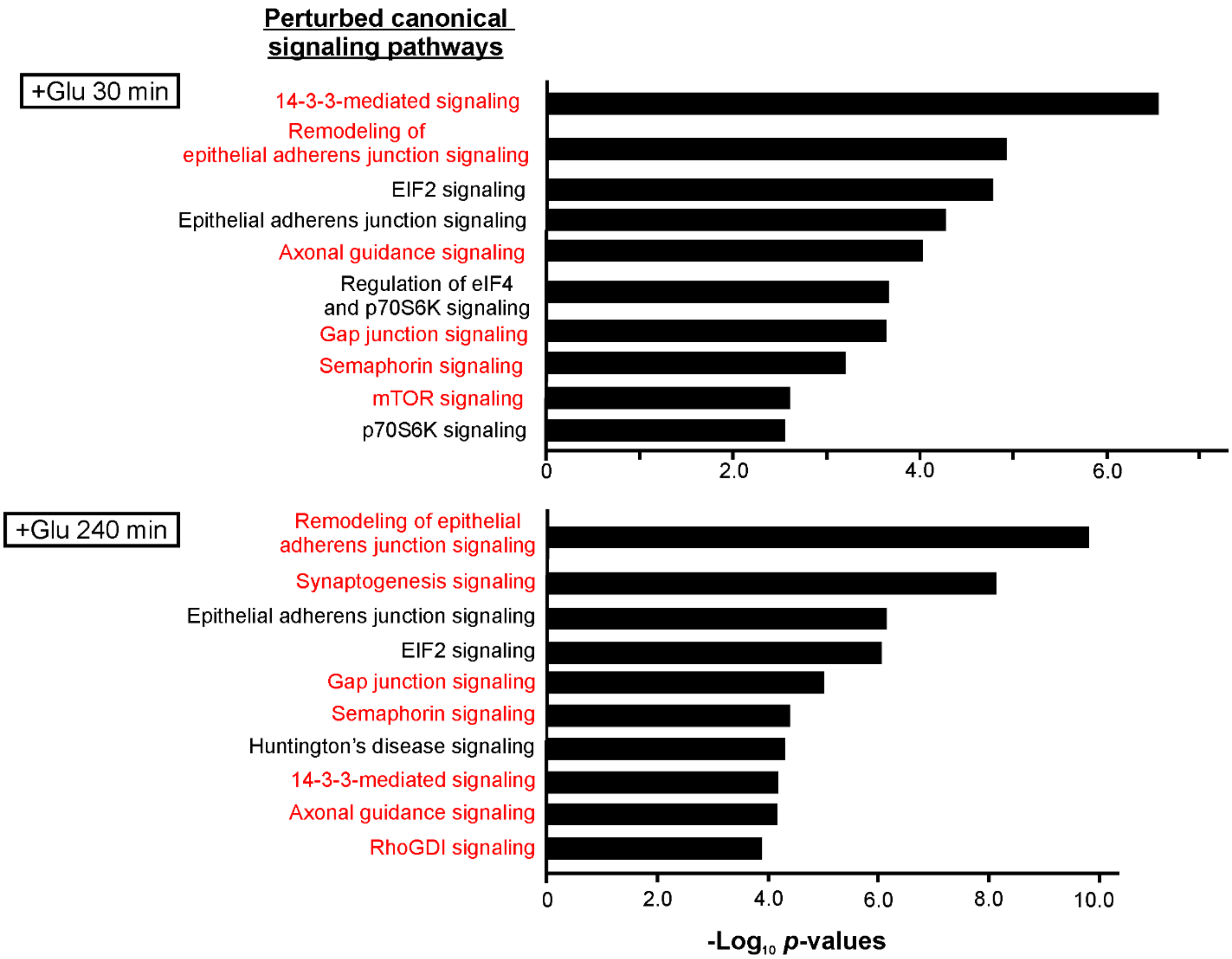


**Figure S9 Canonical signaling pathways and cellular functions in neurons perturbed by glutamate treatment**

Top-ranked perturbed neuronal signaling pathways in response to glutamate treatment revealed by interrogation of the changes in N-terminome of excitotoxic neurons with the Ingenuity Pathway Analysis (IPA) software. The minimum threshold for *p*-value was set to < 0.01. The pathways in red fonts are those involved in synaptic organization and functions.

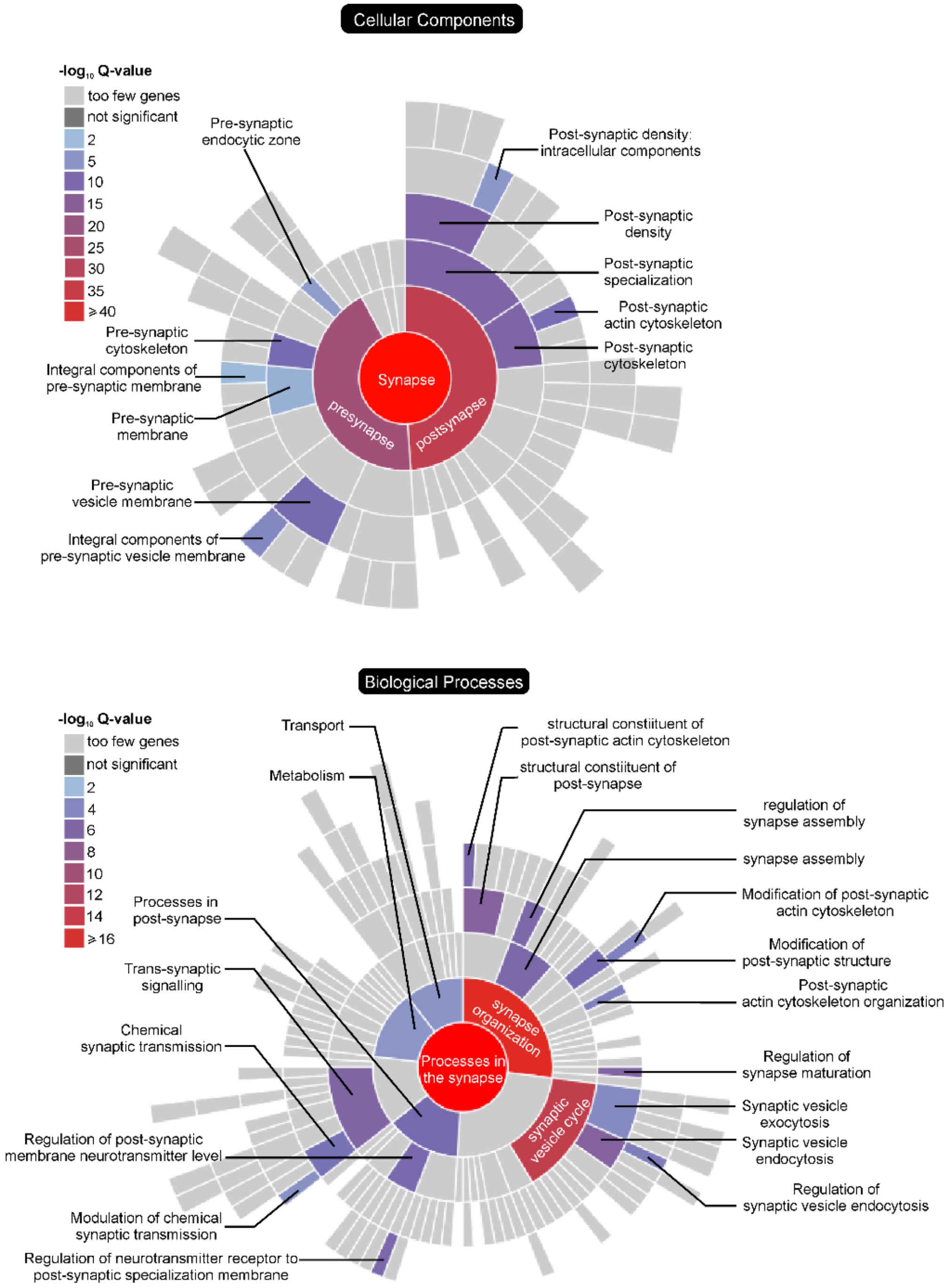


**Figure S10 Sunburst plots depicting the locations (cellular components) and biological functions of neuronal proteins undergoing significantly enhanced proteolytic processing during excitotoxicity**

The heatmaps and their annotations indicate the significance (represented as $-\log_{10}$ enrichment Q values) of enrichment in the indicated locations and biological processes. Neuronal proteins enriched in these locations and participating these biological processes are listed in Table S9 and Figure 2A.

A.

| Cleavage site | M/L Ratio 30 min | M/L Ratio 240 min |
|---|---|---|
| A516-S517 | 2.66 | N.D. |
| S517-S518 | 4.96 | 9.74 |

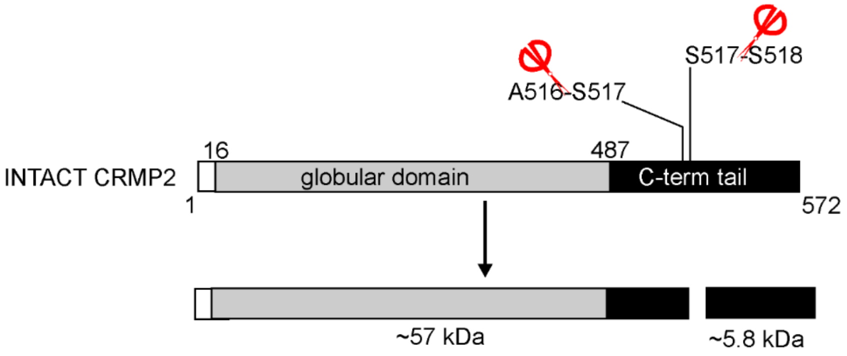


B.

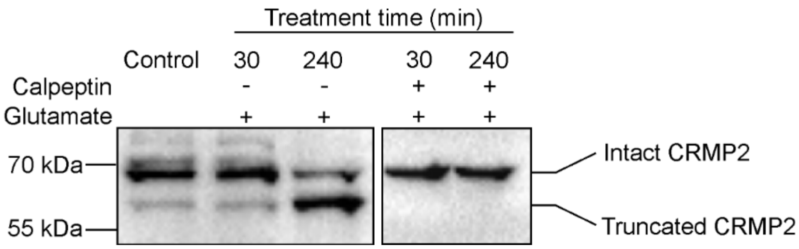


**Figure S11 Calpeptin abolished proteolytic processing of CRMP2 (DPYSL2) to form truncated fragments induced by glutamate over-simulation**

**A.** Functional domains of CRMP2. The sites of proteolytic processing induced by glutamate treatment at\re shown. Inset: the abundance (M/L) ratios of the neo-N-terminal peptides in glutamate-treated neurons versus those in control neurons. These peptides were undetectable in neurons co-treated with glutamate and calpeptin. **B.** Western blots of lysates of neurons treated with glutamate and neurons co-treated with glutamate and calpeptin probed with the anti-CRMP2 antibody. The image of lysates of control neurons and neurons treated with glutamate is also shown in Figure 3B.

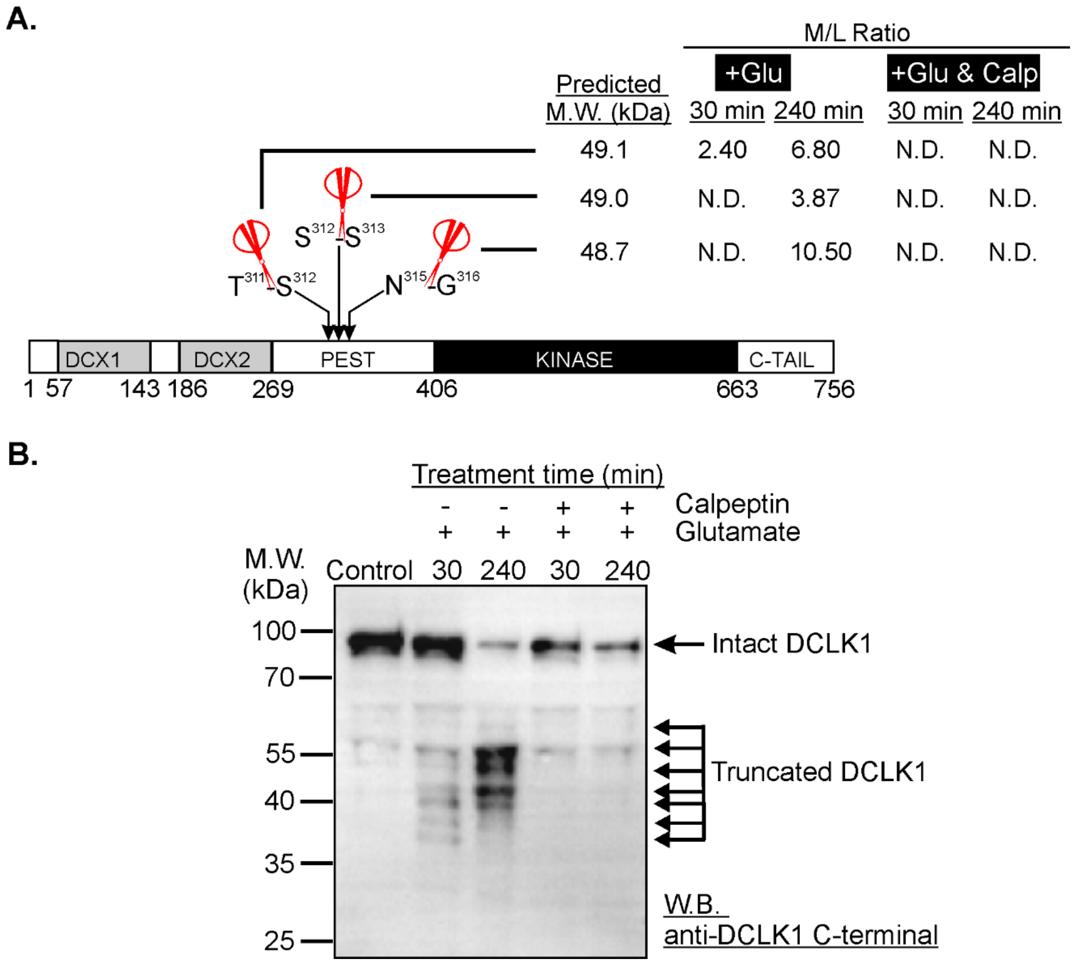


**Figure S12 DCLK1 as a potential substrate of calpains in excitotoxic neurons**

**A.** The cleavage sites in DCLK1 identified by TAILS. Red scissors: cleavage sites of significantly enhanced proteolytic processing during excitotoxicity. DCX1 and DCX2: doublecortin domains 1 and 2. PEST: sequence rich in proline, glutamate, serine and threonine. KINASE: protein kinase domain. C-TAIL: C-terminal terminal. **B.** Western blot of lysates from untreated (Control), glutamate-treated, and glutamate/calpeptin co-treated neurons probed with the anti-C-terminal DCLK1 antibody. Some of the truncated DCLK1 fragments were likely generated by cleavage at sites of significantly enhanced proteolytic processing identified by TAILS shown in panel A.
